# The Evolution of Local Energetic Frustration in Protein Families

**DOI:** 10.1101/2023.01.25.525527

**Authors:** Maria I. Freiberger, Victoria I. Ruiz-Serra, Camila Pontes, Miguel Romero-Durana, Pablo Galaz-Davison, Cesar Ramírez-Sarmiento, Claudio D. Schuster, Marcelo A. Marti, Peter G. Wolynes, Diego U. Ferreiro, R. Gonzalo Parra, Alfonso Valencia

## Abstract

Energetic local frustration offers a biophysical perspective to interpret the effects of sequence variability on protein families. Here we present a methodology to analyze local frustration patterns within protein families that allows us to uncover constraints related to stability and function, and identify differential frustration patterns in families with a common ancestry. We have analyzed these signals in very well studied cases such as PDZ, SH3, *α* and *β* globins and RAS families. Recent advances in protein structure prediction make it possible to analyze a vast majority of the protein space. An automatic and unsupervised proteome-wide analysis on the SARS-CoV-2 virus demonstrates the potential of our approach to enhance our understanding of the natural phenotypic diversity of protein families beyond single protein instances. We have applied our method to modify biophysical properties of natural proteins based on their family properties, as well as perform unsupervised analysis of large datasets to shed light on the physicochemical signatures of poorly characterized proteins such as emergent pathogens.

## Introduction

Families of proteins originate from a common ancestor and develop over evolutionary timescales through various mechanisms of sequence variability at the domain level (1). The effects that these variations have on phenotypic traits such as protein stability and biological function may conflict with one another, restricting evolutionary trajectories in the protein sequence space (2). Multiple sequence alignments (MSAs) of protein families show that there are certain positions under strong evolutionary pressure that have little variability while other positions undergo neutral evolution. The latter ones allow protein sequences to diffuse in sequence space as long as the mutations preserve the structure of their ground states along with their thermodynamic stability and kinetic accessibility while not compromising function (3). Superfamilies, families and subfamilies are terms that have been coined to organize the different levels of sequence, structure and functional similarity as evolution progresses and phylogenetic trees grow. The MSAs of superfamilies show distinctive patterns of differentially conserved residues that modulate the specificity of biological function within different subfamilies. Methods that represent proteins, with their sequences as vectors in a generalized sequence space (4) can identify “Specificity Determining Positions” (SDPs) (5), i.e. positions that are differentially conserved within distinct subfamilies. Nevertheless, interpreting SDPs from sequence alone remains challenging. Energy based, structural approaches become useful, since sequence diverges faster than structure (6), and structural comparison has facilitated to group evolutionarily related protein families into superfamilies, even in the absence of detectable sequence similarity (7). Recent advances in protein structure prediction (8, 9) have made high quality structural models available to most known proteins. Therefore, it becomes possible to comprehensively integrate sequence and structural information to enhance our understanding of the natural phenotypic diversity of protein families.

Sequence variations can be linked with their structural, functional and dynamic consequences using the concept of local energetic frustration (10), derived from the energy landscape theory of protein folding. According to the “Minimal Frustration Principle”, possible strong energetic conflicts between different residues are minimized in the native states of foldable proteins, unlike random heteropolymers. Nevertheless, some conflicts may have been positively selected by evolution due to functional requirements. These conflicting signals have been related to many functional aspects of proteins such as the binding to small ligands or cofactors as well as protein-protein interactions (10), allosterism (11), disorder/order transitions (12) or the existence of fuzzy regions (13). To quantify frustration, the energetic contribution of a particular interaction in the native structure is compared with the distribution of free energy of decoys, i.e. alternative sequence structure pairings. A local frustration index, being defined as the Z-score between the energetics of the native interaction and the distribution of decoys, tells us how well the local energetics has been optimized for folding. A native interaction may be classified as minimally frustrated when most changes in that location destabilize the overall structure relative to the decoys. Conversely, if most local sequence or structural changes would lower the free energy of the system then the interaction will be labeled as highly frustrated (10). Similarly, a single residue frustration index can be obtained when only one residue identity is mutated and all of its interactions are simultaneously evaluated.

Here we explore the concept of local energetic frustration in the evolutionary context of protein families, going beyond single protein instances. The analysis of structural and functional features using the conservation of local frustration permits the identification of common and differential patterns among evolutionary related subfamilies. We demonstrate the usefulness of the evolutionary analysis of frustration in protein families to address several questions. We show how such an analysis can be used to retrieve experimental measurements of physicochemical changes in proteins belonging to the PDZ, SH3 and KRAS families; and study the functional and structural divergence of related protein families such as *α* and *β* globins and the RAS subfamilies. Moreover, we show the general applicability of these ideas by developing an unsupervised strategy to rapidly uncover sequence and energetic constraints in large datasets like the entire SARS-CoV-2 proteome. Finally, we show how such analysis can guide attempts to modify the biophysical behavior of the metamorphic RfaH protein based on its family frustration conservation patterns.

## Results

### 1. The Evolution of Frustration In Protein Families

We have developed a method, called FrustraEvo, to measure the conservation of local energetic frustration over aligned residues or contacts in a protein family. Local energetic frustration measures how well optimized for folding the energy of a given residue-residue interaction is in comparison to the random interactions that would occur within the polypeptidic chain in non-native conformations. Using the Z-score that compares the native energy to the distribution of energies of the decoys, we can classify interactions as being in one of 3 classes: 1) highly frustrated, if most decoys have a lower energy, 2) minimally frustrated, if most decoy possibilities have a higher energy or 3) neutral in an intermediate case scenario (see Methods). There are 3 ways to generate the decoys that give 3 different frustration indices (FI), 2 for pairwise contacts, i.e. mutational and configurational FIs and single residue frustration index (SRFI). Mutational frustration indicates how the frustration changes as a function of the amino acid identities for a given pair of positions. Configurational frustration measures how frustration changes not only relative to amino acid identities but also to the change of conformational state and solvent exposure. Although the two indices are well correlated (10), the mutational FI is somewhat more useful for identifying active or ligand binding sites, while the configurational FI can be employed to analyze protein-protein interactions and conformational changes. The SRFI aggregates the energetic description of all interactions established by a given residue when mutating its identity (see Methods).

Local frustration conservation analysis can be carried out using any of the 3 FIs. For simplicity, we next explain the methodology based on the SRFI although the same analysis can be generalized to the pairwise contacts FIs. Given a Multiple Sequence Alignment (MSA) and the corresponding structures for each sequence contained in it, we can compare the local frustration values from all of the structures at each aligned residue within the MSA. In Fig. 1A we show this mapping for a region of the *α* globin family. Some columns in the MSA show more frustration conservation compared to others. We can quantify the evolutionary significance of such conservation by calculating the Information Content (IC) for each MSA column (see Methods), based on the distribution of local frustration states (FrustIC). The more conserved the frustration state is at a given MSA position, the higher its FrustIC will be and conversely, the lower the FrustIC is the more heterogeneous the distribution of frustration states is at that MSA position. In Fig. 1B we show the SRFI for the human *α* globin protein (FrustratometeR results (14)) as well as the FrustIC values computed from the *α* globins MSA (FrustraEvo results), mapped on top of the human *α* globin structure. While some frustration states at the individual protein are consistent with the ones being conserved at the family level, others are not, reflecting protein specific, perhaps evolutionarily divergent characteristics (Fig. 1B).

**Fig. 1.**
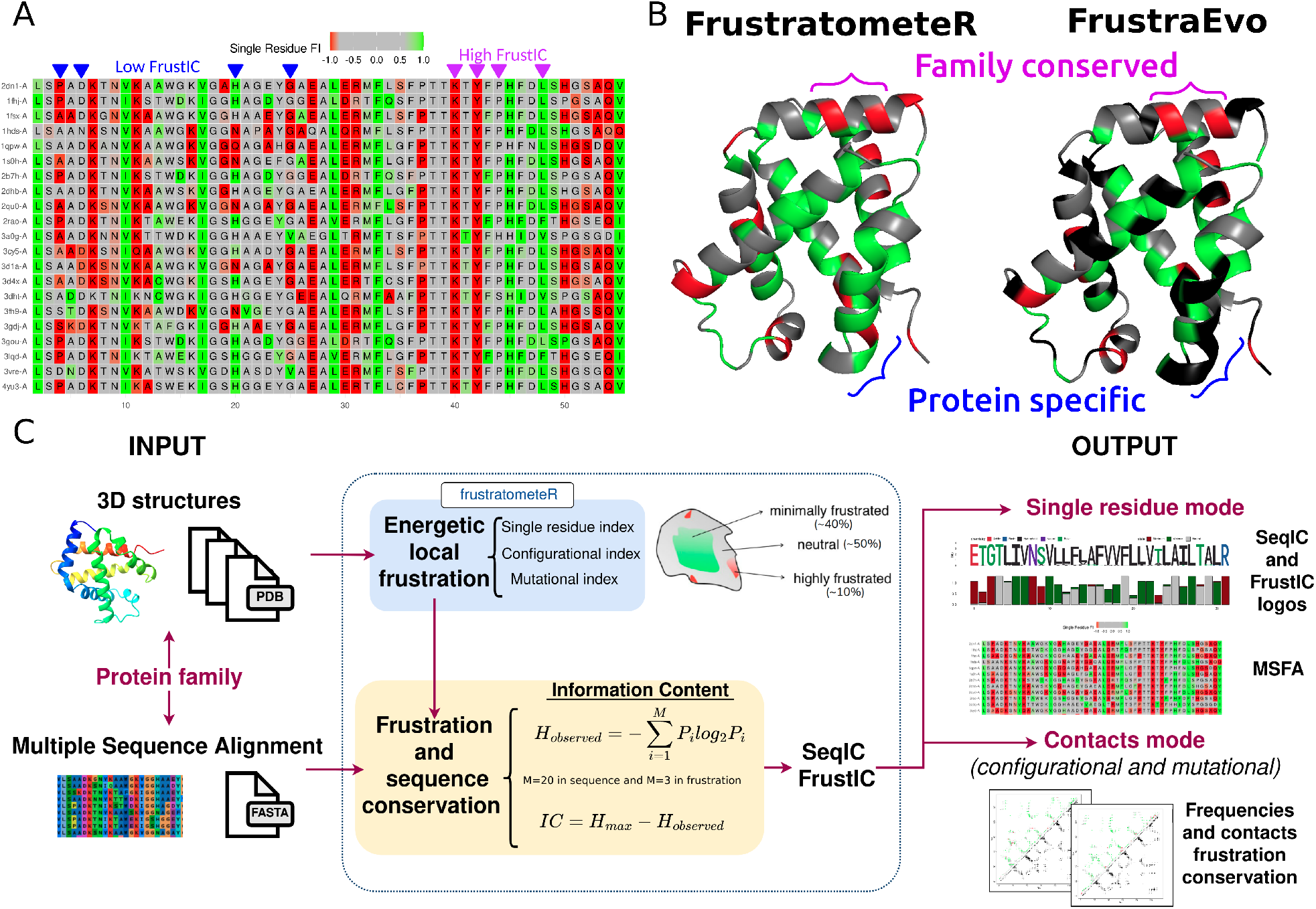
Analysis of frustration in protein families. **A)**. Multiple Sequence Frustration Alignment (MSFA) that consists in the SRFI computed from individual protein structures mapped into the MSA (see Methods). Residues in the MSA are coloured according to their SRFI in the corresponding structures. Magenta asterisks mark frustrationally conserved residues (high FrustIC) and blue ones mark non frustrationally conserved residues (low FrustIC). Minimally frustrated residues are colored in shades of green, neutral in gray and highly frustrated in red. **B)** Comparison between SRFI values as calculated by FrustratometeR (left) and the conservation of frustration states based on their FrustIC values as calculated by FrustraEvo (right) visualized in the same structure (human *α* globin, PDB 2DN1, chain A). In the FrustratometeR representation residues are coloured according to their frustration states. In the FrustraEvo representation residues with FrstIC*>*0.5 are coloured according to their most informative frustration state while residues with FrstIC*≤*0.5 are coloured in black. **C)** Overview of the FrustraEvo workflow to analyze a single protein family.

A schematic view of FrustraEvo’s workflow is shown in Fig. 1C (see Methods for details).

### 2. Evolutionary Analysis of Frustration Unveils Stability Constraints Within Protein Folds

To investigate the evolutionary role of frustration in stability and function we analyze three specific family cases for which double-deep protein fragment complementation (ddPCA) experiments have been performed and are available. This experimental method has been used to quantify the effects of amino acid variation on protein stability (abundance) and function (binding) (15, 16). For each position of the C-terminal SH3 domain of the human growth factor receptor-bound protein 2 (GRB2-SH3), the third PDZ domain of the adaptor protein PSD95 (PSD95-PDZ3) and the GTPase KRas (KRAS), the experiments have produced two scores (ddPCA phenotype scores), corresponding to the effect of variations on stability and function. The abundance and binding ddPCA scores are largely correlated for SH3 and PDZ while this correlation is weaker for the KRAS case.

We automatically retrieved homologous proteins for the 3 studied proteins, although for KRAS we also analyzed a highly curated dataset from Rojas et al (17) (see Methods). We analyzed the relationship between the calculated SeqIC and FrustIC values for individual MSA positions with the ddPCA experimental abundance scores derived for the individual proteins. We found that FrustIC is a good predictor of the experimental ddPCA phenotype scores for stability and function for SH3 (r=-0.79, p-value=9.5e-05) and PDZ (r=-0.82, p-value=7.6e-08) (Fig. 2C, 2E and S2A and S2D). The SeqIC score is also correlated with the ddPCA phenotypes although with a slightly lower Pearson correlation coefficient (r=-0.63, p-value=2.5e-07 and r=-0.69, p-value=6e-13, respectively) (Fig. 2B, 2D, S2B, S2E). Furthermore, SeqIC and FrustIC are both correlated to each other in both proteins (Fig. S2C and S2F). On the other hand, KRAS shows no significant correlation between the ddPCA scores and SeqIC (Fig. 2F) but has a significant and moderate correlation with FrustIC for the minimally frustrated and conserved residues (r=-0.47, p-value=0.00012) (Fig. 2G). The correlation between SeqIC and FrustIC is also weaker (Fig. S2I). Most residues with FrustIC values close to the theoretical maximum (*log*_2_(3)=1.58, see Methods) are located in the hydrophobic core of the protein (Fig. S1C). Because they can vary between the hydrophobic types of amino acids without affecting stability too much, their FrustIC values tend to better correlate with abundance ddPCA score than the SeqIC values (Fig. S1B). Relationships between SeqIC, FrustIC and the ddPCA binding score are shown in Fig. S2.

**Fig. 2.**
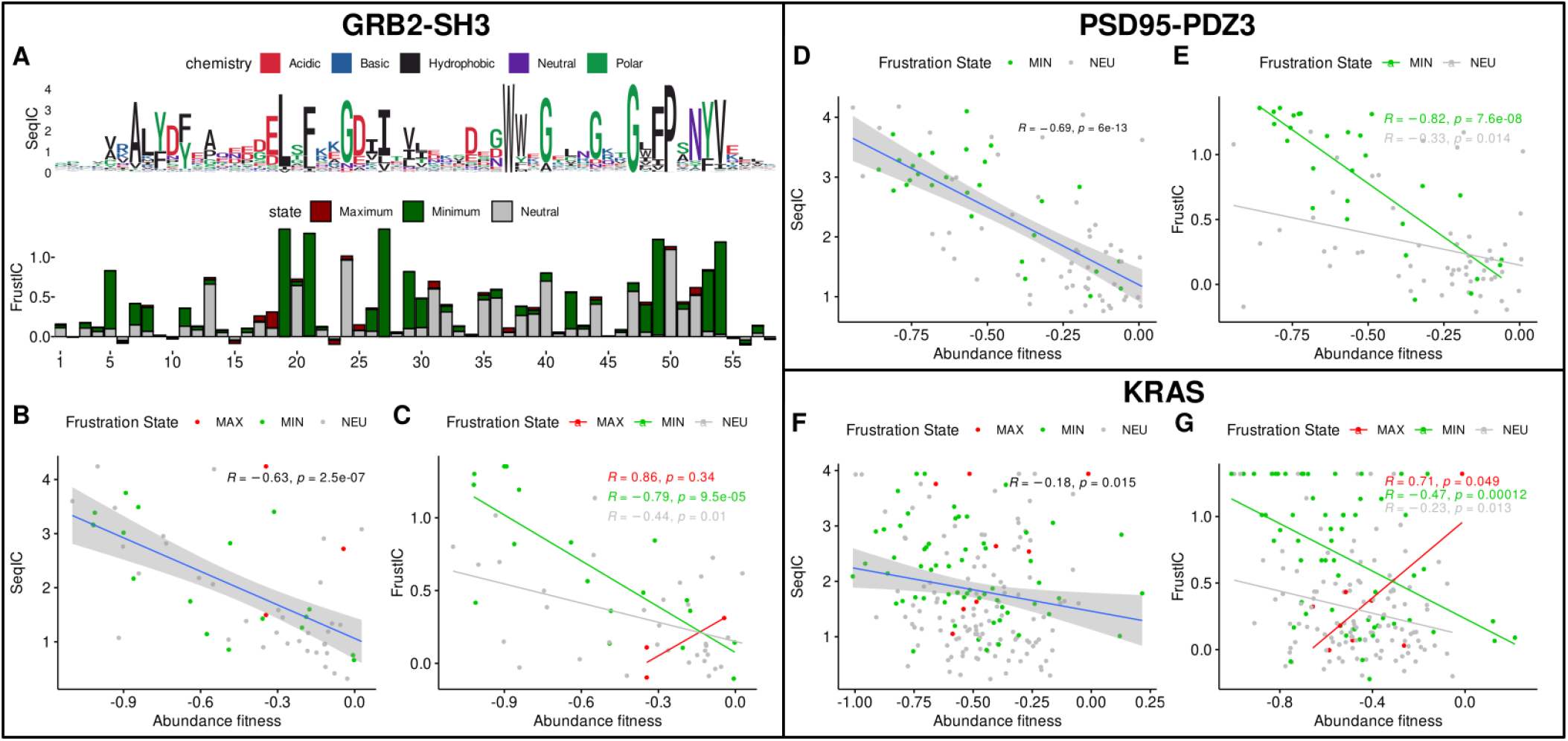
FrustIC correlates with experimentally measured protein stability changes. **A)** Sequence and Frustration logo plots showing SeqIC and FrustIC values per MSA column respectively for GRB2-SH3. The numbering of the plot corresponds to the sequence of reference (chain A from PDB 2VWF in the case of GRB2-SH3). Positions containing a gap in the sequence of reference are not considered in the plot. **B-G)** Correlation between ddPCA abundance scores vs SeqIC and FrustIC for GRB2-SH3, **B-C)** PSD95-PDZ3, **(D-E)** and KRAS **F-G)**.

Both SH3 (Fig. 2A) and PDZ domains (Fig. S1A) do not have highly frustrated conserved positions (FrstIC*>*0.5). As these are protein-protein interactors with no localized function, highly frustrated residues are not aligned in the family MSAs and hence their signal gets averaged out. In contrast KRAS has one highly frustrated conserved position (FrustIC*>*0.5, Fig. S1B), Lys117 (KRAS numbering), which is one of the seven conserved residues that interact with the nucleotide substrate (18). In addition, Lys117 has a very high ddPCA value. This means that most mutations in that residue improve the protein stability, reinforcing that functional and stability signals often conflict with each other at that locus. The results for KRAS that have been discussed correspond to the curated dataset from Rojas et al. We further repeated the analysis using the automatic retrieval of homologous proteins (see Methods) and found that the same trends are recovered although the correlation between FrustIC and the abundance ddPCA score is weaker (FigS3B). This might be because of two reasons: 1) this dataset contains 1354 proteins vs the 36 contained in the Rojas et al. one and 2) the homologous relationship in the Rojas et al. was defined by phylogenetic studies while in the other it was assumed after performing Blast and simple quality filters (see Methods). This highlights that although our strategy can retrieve meaningful results from automatically generated datasets, highly curated ones will perform better.

Altogether, these examples show that frustration conservation analysis can predict residues that are related to protein stability with experimental correlation. For protein-protein interactors, like PDZ and SH3, SeqIC alone predicts much of the experimental ddPCA scores although FrustIC at the minimally frustrated residues is a somewhat better predictor. For other architectures with more localized function, like KRAS, only FrustIC significantly correlates with the experimental scores. Frustration conservation analysis can further identify residues that are important for function, like Lys117, which in addition shows a stabilizing effect when mutated, as measured by ddPCA.

### 3. Differential frustration conservation patterns reveal lineage specific functional adaptations within protein superfamilies

We further investigated the link between sequence divergence and the divergence of local energetic frustration by analyzing variability among evolutionarily related, but distinct, protein families. We analyzed the common and differential frustration conservation patterns for two very well studied examples; first comparing the *α* and *β* globins, which are parts of the hemoglobin biological unit, and then examining the human RAS superfamily. The globins are relatively closely related with each other while the RAS superfamily has experienced wide sequence divergence.

The *α* and *β* globin subfamilies have a common origin, but despite their very similar structures they have different and well-studied functions, as part of the hemoglobin *α*2*β*2 tetramer (19). We used a non redundant set of experimental structures that correspond to 21 mammalian hemoglobins (see Methods). Fig. 3A shows the frustration conservation patterns, based on the SRFI, for the *α* and *β* lineages grouped into a single dataset (*α*/*β* dataset). Frustration level is mostly conserved (FrustIC*>*0.5) at minimally frustrated positions (n=35, mean FrustIC=1.02) and at neutral positions (n=34, mean FrustIC=0.85). Only 3 positions are highly frustrated (mean FrustIC=0.72).

**Fig. 3.**
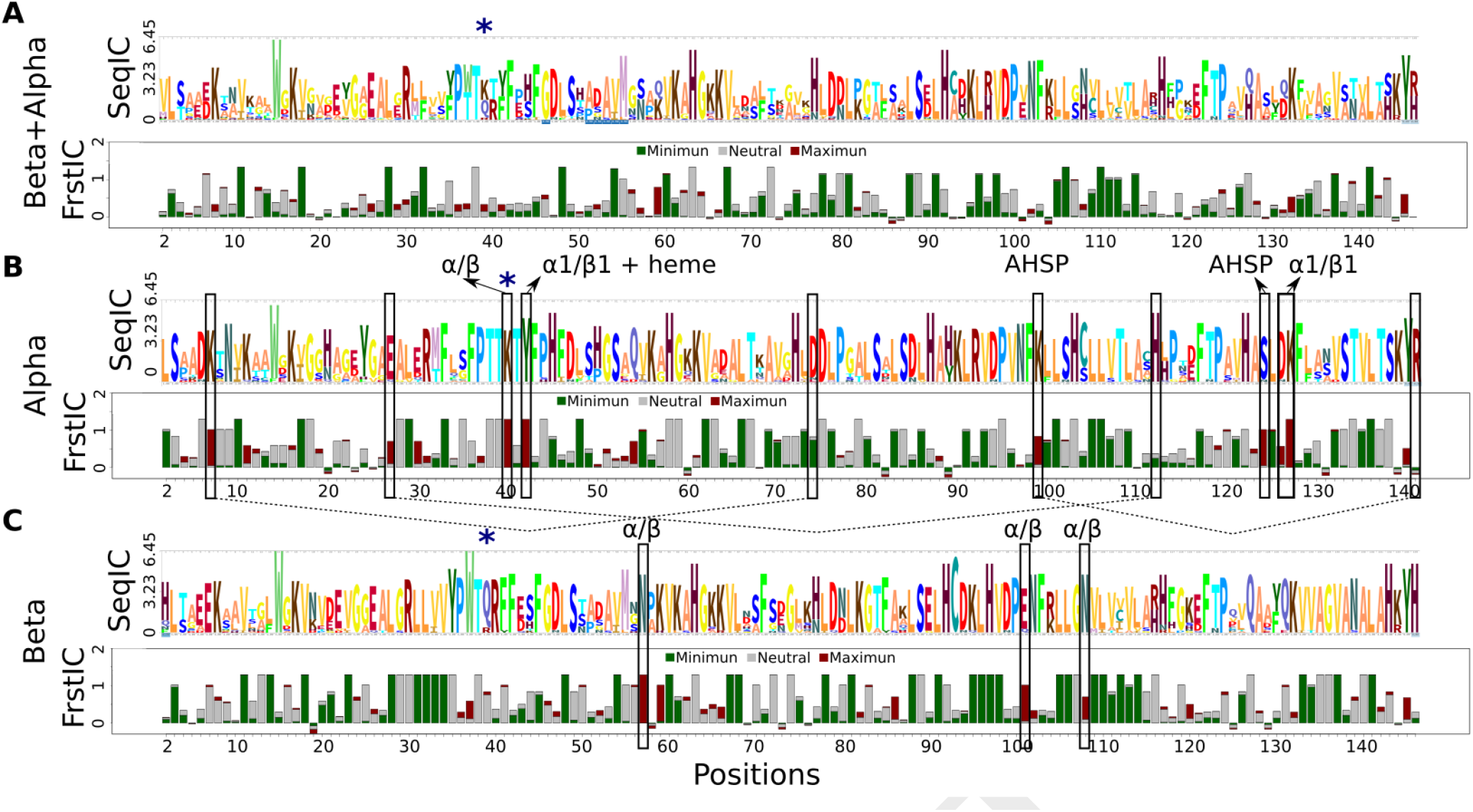
Differential frustration conservation patterns unmask functional constraints in the hemoglobin subunits. FrustraEvo results based on the SRFI for **(A)** *α*/*β*-globins, **(B)** only *α* and **(C)** only *β*. Rectangles denote functionally relevant positions explained in more detail in Table S1. In blue asterisks we marked position 39 in the *α*/*β* MSA, that corresponds to a highly frustrated Lys40*α* but to a neutral Glu39*β*.

Some positions in the *α*/*β* MSA show changes in their amino acid identity that result in differential frustration conservation signals when the two lineages are analyzed separately. An example of this is position 39 in the *α*/*β* MSA, that corresponds to a highly frustrated Lys in the *α* lineage (Lys40*α*) (Fig. 3B) but to a neutral Glu in the *β* lineage (Glu39*β*) (Fig. 3C). This suggests that this position corresponds to a functional adaptation that occurred after the divergence of the two families and that there are more functional constraints associated to this position in the *α* family compared to the same position in the *β* family. This residue represents an example of a SDP (5) between sub-clusters in a MSA, to which evolutionary frustration analysis can provide a functional explanation based on the FrustIC values in each family, as we will extend later for other example.

In total, there are 12 highly frustrated positions in the *α* globins (mean FrustIC=0.87, Fig. 3B) and 8 in *β* globins (mean FrustIC=0.88, Fig. 3C). Most of these positions are frustrated in only one of the two families. These loci have been described as being involved in protein-protein interactions within the tetrameric structure of Hemoglobin or occur between the *α*-globins and the *α*Hb-stabilizing protein (AHSP), a chaperone that prevents *α*-globin toxicity when isolated (20, 21) (Table S1). Several other highly frustrated and conserved residues (FrustIC*>*0.5) in *α*-globins correspond to residues that are part of intra chain salt bridges (22) which are critical for allostery and the Bohr effect as explained by Perutz (23) (Fig. 3B, Table S1).

Comparing superfamilies that have a distribution of broader evolutionary distances among their members proves more challenging than for the case of *α* and *β* globins. The human RAS superfamily comprises a broad range of proteins that despite their sequence and functional diversification, share both a common structural framework and enzymatic activity related to the production of GDP by the hydrolysis of GTP (17). These proteins exist in two conformational states: an active GTP-bound state capable of binding to effectors to transduce signals and an inactive GDP-bound state. There are 5 RAS subfamilies, i.e. RAS, RHO, RAB, ARF and RAN, that bind to different effectors. The conserved GTP-binding site consists of 5 motifs, named G1 to G5, that contain the core residues essential for the GTPase activity and its associated conformational changes. These motifs have been shown to harbor different SDPs linked to family-specific functionalities (17).

We retrieved all paralogs within the RAS superfamily in humans (17) and calculated their frustration conservation patterns (see Methods). The RAN family was omitted from our analysis as it contained only one sequence. As a consequence of the high energetic variability within each family, the frustration conservation analysis using the SRFI analysis shows little conservation for the G1-G5 motifs (Fig. S4) with only a few minimally frustrated and neutral positions being frustrationally conserved (FrstIC*>*0.5) across all families. Consequently, we also explored frustration conservation at the level of residue-residue contacts based on the mutational FI. This index was used instead of the configurational one as we expect functional signals to be more dependent on the amino acids identity than the conformational state of the protein. When mapped onto the reference structures for each RAS family, highly frustrated interactions are localized around the G1-G5 motifs whereas the rest of the fold is mainly enriched in minimally frustrated interactions (Fig. 4A). To enhance interpretability, we represented the interactions involving only residues that belong to the G1-G5 motifs or that are SDPs in each subfamily as graphs (Fig. 4B). Despite the large evolutionary distances, a network of highly frustrated interactions involving mainly Lys16 (G1), Asp57 (G3) and Lys117 (G4) is present in all four subfamilies. These interactions involve residues that interact with the GTP or the Mg ion as well as some residues that interact with other protein partners (Fig. S5). RHO is the most divergent family as it has an extra helix next to G4, after which there appears Pro138 (Pro124 in RAS numbering) that participates in many highly frustrated interactions with Lys117 and Lys118. These interactions are part of a different highly frustrated interaction network from the one containing Lys16 and Asp57.

**Fig. 4.**
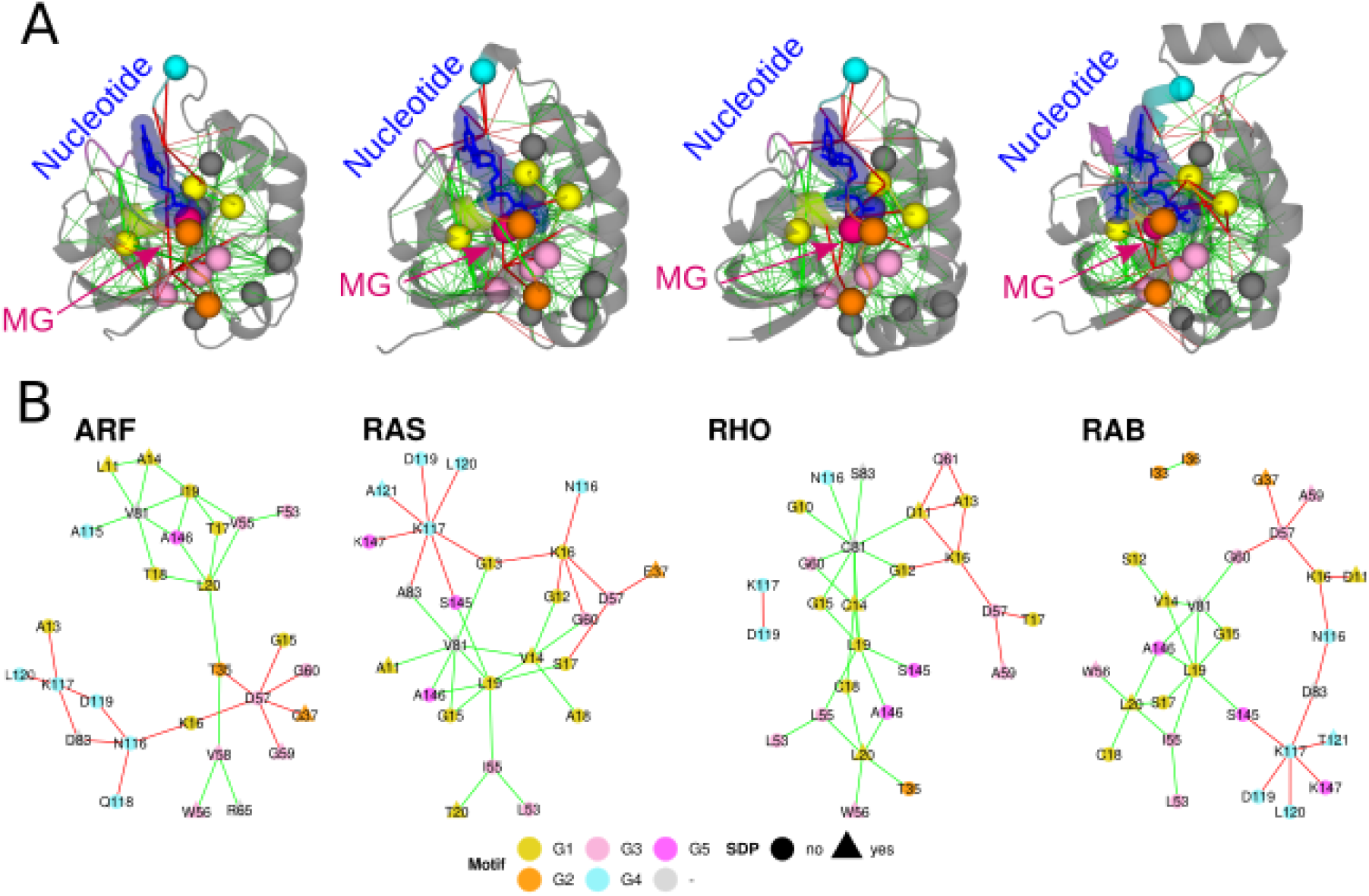
Frustration conservation patterns unmask strongly conserved constraints across the Human RAS superfamily. **A)** Conserved highly frustrated (red lines) and minimally frustrated (green lines) interactions (FrustIC*>*0.5) according to the mutational FI. *C*_*α*_ of SDPs are in globular shape. G1 motif is in yellow, G2 orange, G3 light pink, G4 cyan and G5 in magenta. Structures shown correspond to PDBs 7MGE (ARF), 3TKL (RAB), 121P (RAS) and 6BCB (RHO). **B)** Networks representing conserved highly frustrated (red lines) or minimally frustrated (green lines) interactions (FrustIC*>*0.5) between residues within the G1-G5 motifs (circular shape) or SDPs (triangular shape) in at least 50% of the structures of each subfamily. Nodes with triangular shape correspond to SDPs outside the G1-G5 motifs.

Beyond similarities, there are energetic differences between the different subfamilies which seem to be related to SDPs (marked with asterisks in Fig. S4). As an example, SDP 83 (RAS numbering) shows family specific requirements; being neutral or without local frustration conservation at the single residue level but with differentially conserved frustration in the interactions with other residues. This SDP is a highly conserved Asp with highly frustrated interactions (Fig. 4B) in ARF and RAB. In contrast, SDP 83 is a conserved Ser in RHO that establishes minimally frustrated interactions. The SDP identity is not conserved in RAS, having a mixture of both highly frustrated and minimally frustrated interactions (the latter with Val81, another SDP). Other SDPs, e.g. 20, 56 and 81 (RAS numbering) are minimally frustrated with variable identities within the hydrophobic and polar group of amino acids. The change in identity within the SDPs that interact with each other seems to be evolutionarily compensated in each subfamily, possibly finely tuning specific interactions with other residues.

Frustration conservation analysis allows one to interpret sequence diversity among evolutionarily related families and to link this diversity with functional adaptations within divergent protein lineages. In some cases, as for the globins, frustration conservation analysis using the SRFI is sufficient to uncover these functional and stability signals while in more divergent examples, conservation analysis using the pairwise contacts FIs is more enlightening.

### 4. Large Scale Application Of Frustration Conservation Analysis In Coronaviruses

A valuable use of analyzing the conservation of frustration is to provide insights about proteins that are still poorly characterized in the laboratory, such as those from novel, emergent pathogens. To illustrate this application, we have automatised the steps of generating MSAs, clustering the resulting subfamilies based on the SDP methodology (5), and structural models prediction by AlphaFold2; and combined these steps with the use of FrustraEvo using the SRFI. We applied this workflow to analyze the full SARS-CoV-2 proteome in the context of the entire coronaviruses phylogeny (see Methods; data available in Zenodo, see Materials). Here, we present a descriptive overview of the results for the full proteome and provide a more detailed analysis for one of the coronavirus proteins, the papain-like protease (PLpro), essential for viral replication (24, 25).

We have used our pipeline to process 29 SARS-CoV-2 proteins or protein domains (see Methods) from which 22 of them passed our quality filters (more than 10 sequences and average pLDDT over aligned positions of the cluster ≥ 80; see Methods, Fig. S6). For each cluster we compared sequence and frustration conservation by calculating the mean SeqIC and FrustIC per position and per cluster. In Fig. 5A we observe a significant positive correlation between these values (r=0.69, pvalue=2.6e-14). Some protein families like cluster 5 in the C-terminal domain of the Nucleoprotein (N_CTerm) deviate from the correlation. These have a lower FrustIC than expected (Fig. S7). Close inspection of this case shows that this family has regions with low pLDDT scores (Fig. S8A) regardless of the median for the entire protein being pLDDT ≥ 80, which is known to correlate with flexible regions (26). Hence the conformation of the region and its frustration values can be heterogeneous and therefore yield lower FrustIC than expected given the sequence diversity in the family. At the same time, however, other factors such as the predicted amount of disordered residues (Fig. S8B) or the MSA phylogenetic diversity (Fig. S8C) can also influence the FrustIC and SeqIC relationship. On the other side of the spectrum, we have cases like cluster 2 in the non-structural protein 13 (nsp13) (Fig. S7) for which the analysis of frustration conservation explains much more than expected from the overall correlation with sequence, because at some loci different amino acids can have similar frustration states, e.g, hydrophobic residues in the core of the structure (Fig. S9). Finally, when assessing full protein families we have found that some proteins, e.g. the C-terminal of the SARS-Unique Domain (SUD_Cterm) of nsp3 (3 subfamilies, FrustIC sd=0.042) or the Envelope (E) protein (2 subfamilies, FrustIC sd=0.046) are very homogeneous in terms of their average FrustIC values across subfamilies while others,e.g. N_Cterm (5 subfamilies, FrustIC sd=0.217), N_Nterm (6 subfamilies, FrustIC sd=0.20) or nsp8 (6 subfamilies, FrustIC sd=0.176), show a large amount of energetic variability across them. Once again, the quality of the models, the phylogenetic variability and flexibility of protein regions are factors that might influence these observations but no clear trend was identified (Fig. S8). Further information for all the families can be found in Table S2 and Table S3.

**Fig. 5.**
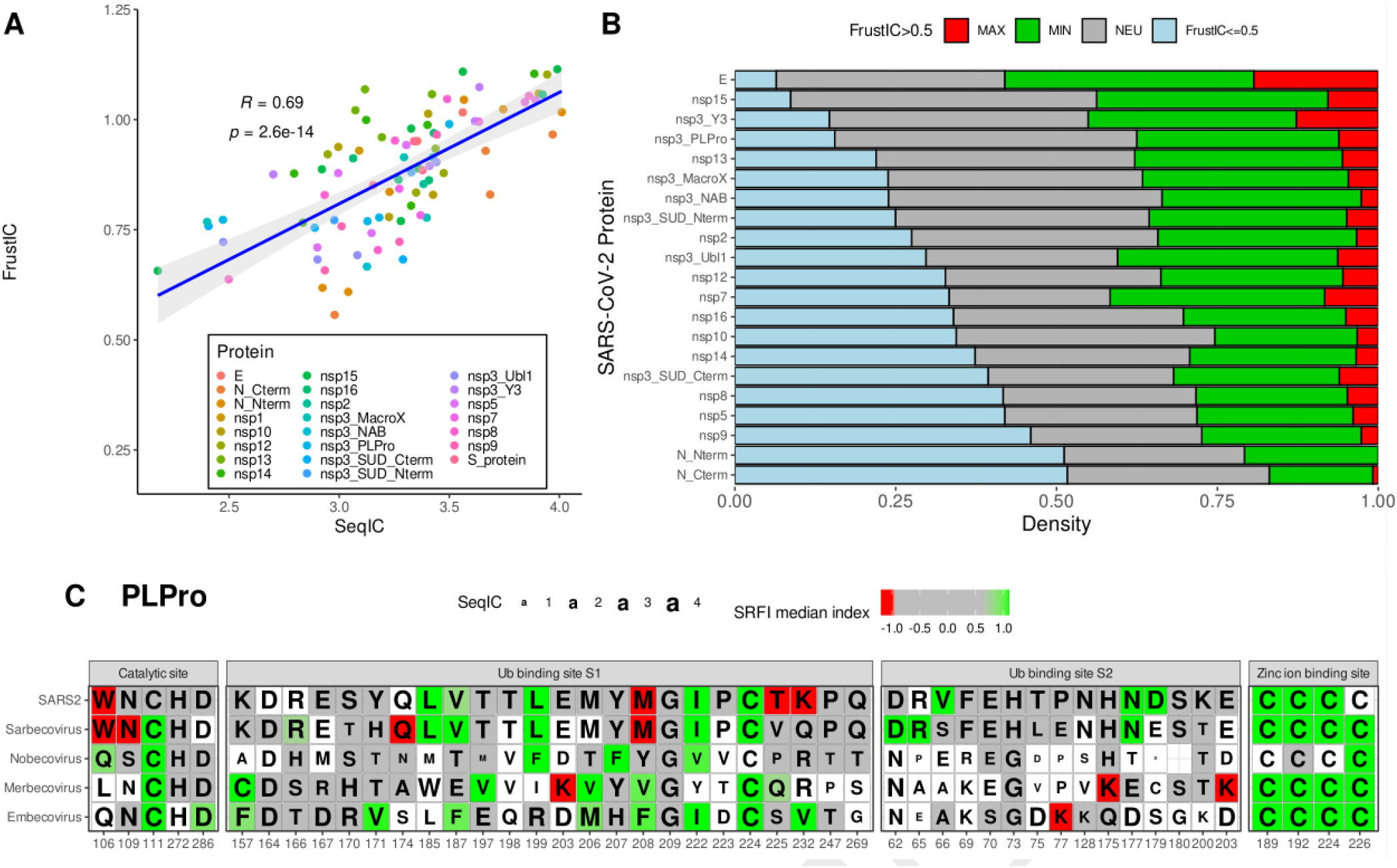
Large scale application of frustration conservation analysis in Coronaviruses. **A)** Correlation plot showing mean FrustIC vs mean SeqIC per S3Det cluster computed for Coronavirus proteins (see Methods). **B)** Distribution of frustrationally conserved residues (FrustIC*>*0.5) for each S3Det cluster containing the corresponding SARS-CoV-2 protein. We considered frustration conservation when FrustIC*>*0.5. The proportion of each protein is normalized by their length (see Table S2). **C)** Multiple sequence/frustration alignment showing FrustraEvo results for selected functional domains in PLpro. Cells that are colored correspond to FrustIC*>*0.5 while white cells mean that FrustIC*≤*0.5. Color of the cells represent the median SRFI value computed with FrustratometeR (see methods for state definition). The amino acid identities correspond to the consensus sequence and the size of the letter is proportional to SeqIC.

As we were interested in using this type of analysis to detect evolutionary constraints in SARS-CoV-2, we next focused on analyzing the Sarbecovirus subfamily that contains the proteins of this virus. In Fig. 5B we show the proportions for frustrationally conserved (FrustIC*>*0.5, either MIN, NEU or MAX frustrated) and non-conserved (FrustIC ≤ 0.5) residues. This type of analysis allows us to rapidly sort proteins according to their frustration conservation level, which ultimately relates to there being different degrees of selective pressure on different proteins of the virus. The proportion of frustrationally conserved residues across proteins is heterogeneous (93.6% in the case of the E protein and up to 48.3% for the N_CTerm domain) reflecting the fact that some proteins are much more constrained by their family features than others. From those that are frustrationally conserved, the majority of residues belong to the neutral (mean=0.35, sd=0.06) or minimally frustrated (mean=0.29, sd=0.06) classes, as expected. An interesting aspect to quantify is how much consistency there is between the Sarbecovirus family and the SARS-CoV-2 proteins. This would let us know to what extent the family restrictions are still present in the virus, whether the restrictions are of a different nature, i.e. stability vs. function, or whether there are novel constraints in the SARS-CoV-2 that were not present in the family. We found 332 residues conserved in the highly frustrated state within the proteome of the SARS-CoV-2 family (Table S4) from which 301 residues were also conserved and highly frustrated in SARS-CoV-2 itself. This suggests that the functional signals, related to the presence of highly frustrated interactions in those regions, are coherent both at the family and at the SARS-CoV-2 levels. We found several residues where the frustration conservation state differs between SARS-CoV-2 and its family. There are 62 residues that are conserved (FrustIC*>*0.5) in the neutral or minimally frustrated state at the family level but conserved in a highly frustrated state in SARS-CoV-2, suggesting recent gain of function events (Table S5). While the majority of these residues are located in proteins that are well studied (Spike n=18, nsp5 n=6), it is noticeable that other less characterized ones contain many of these type of residues as well (nsp2 n=9, nsp3 domains n=9), defining interesting positions for their study (Table S5). In addition, there are 345 positions that are conserved and highly frustrated in SARS-CoV-2 but that are frustrationally conserved at the family level (Table S4).

To better exemplify the impact of this evolutionary style of analysis we describe detailed results for one specific example: the PLpro domain. PLPro catalyzes the proteolysis of the viral polyproteins 23 using a catalytic triad (Cys111, His272, and Asp286) (25) along with two extra residues Trp106 and Asn109 (27). Moreover, PLPro interacts with at least two host proteins, ubiquitin-like interferon-stimulated gene 15 protein (ISG15) and ubiquitin (Ub), to evade or at least hamper the host immune response (28). SARS-CoV-2 PLpro homologous proteins were automatically divided into 4 sub-families that reflect the Betacoronavirus subgenera classification, i.e.: Sarbecovirus (n=31), Nobecovirus (n=11), Merbecovirus (n=35) and Embecovirus (n=45) (Table S3). Additionally, we have manually analyzed a fifth group that only contains experimental SARS-CoV-2 PLpro structures (n=29) to quantify frustration conservation specific to this virus. We compared frustration conservation between the 4 subfamilies to disclose functional diversity related to differential infectivity or virulence (Fig. 5C). At the catalytic site, we observe that the SDP Trp106, which facilitates catalysis (28) by stabilizing the catalytic triad, is highly frustrated only in the Sarbecoviruses group. In the case of Merbecovirus, there is a conserved Leu at position 106, which is reported to make catalysis less efficient (29). When in MERS a Trp is introduced in the analogous position, catalysis is enhanced, suggesting that increased local frustration may be related to the improvement of the catalytic function. In contrast, the catalytic residue Cys111 is minimally frustrated and conserved in all sublineages, reflecting its functional importance to the full phylogeny. In the SARS-CoV-2 set of structures, this position appears as neutral, due to the occurrence of a subgroup that contains the Cys111Ser mutation, which introduces local instabilities. Likewise, the four cysteines that coordinate the binding of an ion of Zinc (Cys189, Cys192, Cys224 and Cys226) (30) are all minimally frustrated and conserved in most of the coronavirus lineages, suggesting a strong stability requirement in that region (Fig. 5C). In fact, the PLpro is unable to function without the binding of this element (30).

Additionally, we observe that S1 and S2, the binding sites to ISG15 and Ub host proteins, are differentially conserved when comparing Sarbecoviruses and the other 3 Betacoronavirus subfamilies (Fig. 5C). The SARS-CoV-2 S1 site contains more highly frustrated residues while the S2 site contains more minimally frustrated residues than do the other viruses. This could explain the differential preference that PLpro has for binding to ISG15 or Ub in SARS-CoV and SARS-CoV-2 (31). For instance, there are positions within the S1 region that become frustrated only at the SARS-CoV-2 level. That is the case for positions 225 and 232 (SARS-CoV-2 numbering) (Fig. 5C) corresponding to neutral Val and Gln in Sarbecovirus but to highly frustrated Thr and Lys in SARS-CoV-2. It has been experimentally shown that these sequence changes affect Ub association, explaining the differential activity on Ub substrates but not on ISG15 (28). Thr 225 is only present in SARS-CoV-2 and in RaTG13 (Fig. S10), the latter being a likely bat progenitor of the COVID-19 virus (32). Moreover the bat-derived viral strains, Rc-o319 and bat-SL-CoVZXC21, contain a Met in that same position that is even more frustrated (Fig. S10). This may point to a position of concern for novel human-infecting variants that could also acquire this change of identity. Glu232 is unique to SARS-CoV-2 within the Sarbecovirus family.

We have shown that the automatic definition of protein families, when combined with frustration conservation analysis, can be proficiently used to detect energetic frustration signatures at different phylogenetic levels in large sets of evolutionarily related proteins. Analyzing the conservation of frustration provides a valuable approach to “blindly” study stability and functional constraints in protein superfamilies and should become a particularly useful resource in future scenarios of emergent pathogens or in the case of understudied protein families.

### 5. Perturbing Conserved Frustration Patterns To Engineer Conformational Changes

Conserved frustration patterns can be used in an applied context to redesign specific functional constraints in specific proteins based on the restrictions observed in their families. In this example, we show how to use frustration conservation to better understand biophysical mechanisms and to engineer, *in silico*, a conformational change in RfaH, a bacterial elongation factor.

The C-terminal domain (CTD) of RfaH undergoes a large, reversible structural rearrangement from an *α*-hairpin (*α*CTD) into a *β*-barrel (*β*CTD) upon interacting with RNA polymerase and specific DNA elements called ops (33). In the absence of ops elements, RfaH *α*CTD is autoinhibited by being bound to the N-terminal domain (NTD) hence, preventing the correct interactions with the RNA polymerase. On the other hand NusG, a non-metamorphic paralog from which RfaH is believed to have originated via gene duplication divergence, only exists in its *β*CTD fold (34). As the RfaH capacity to undergo metamorphosis seems to have occurred after its divergence from NusG, we performed frustration conservation analysis to find the constraints that are present in the RfaH subfamily when in its autoinhibited *α*CTD conformation and explore which perturbations would facilitate its transition into the *β*CTD fold.

We retrieved a set of non-redundant, evolutionarily related RfaH protein sequences (see Methods), predicted their structures with AlphaFold2 (see Methods) and computed their frustration conservation patterns using both the SRFI and the mutational and configurational pairwise contacts FIs. The analysis based on the SRFI identified only two residues that are consistently frustrated across all family members (predominantly red positions in Fig. S11 with FrustIC*>*0.5). Their structural location, far away from the metamorphic domain (residues 110-162), suggests that they do not play a role in RfaH fold-switching. In contrast, the analysis of frustration conservation results based on the contact pair-based configurational and mutational FIs revealed many minimally frustrated and conserved contacts located at the interdomain interface between the *α*CTD and NTD domains (Fig. 6A). This is consistent with the stabilization via interdomain interactions (35, 36) which trigger the RfaH fold-switch towards the *β*CTD conformation (37, 38) when disrupted. We selected 9 interdomain residues (L6, F51, L96, F126, I129, L141, L142, L145, I146) according to their contribution to the interdomain interface stabilization between RfaH NTD and *α*CTD (see Methods) and used FrustratometeR to predict the changes in frustration when individually mutating them to all other 19 amino acids (Fig. S12). Fig. 6B illustrates how the local frustration changes upon mutation between F51 and all the residues with which it interacts. Most of the 21 contacts formed by F51 (Fig. 6B, blue letters) are minimally frustrated, with the exception of 5 that are neutral. Overall, some mutations yield similar frustration values across all contacts (e.g. F51M), while others switch from minimally frustrated interactions to neutral or highly frustrated (e.g. F51K). The same effect is observed when repeating the analysis for the remaining 8 interdomain residues (Fig. S12), leading to the identification of two types of mutations: 1) “Similar Frustration Mutations” (SFMs), which would maintain the stabilizing nature of the native amino acid identities (L6I, F51M, L96W, F126W, I129V, L141V, L142V, L145M, I146V) and 2) “Highly Frustrated Mutations” (HFMs), which would maximize the local frustration index with their neighboring residues (L6D, F51K, L96K, F126N, I129E, L141D, L142K, L145E, I146D). We generated two E. coli RfaH mutant sequences containing all SFMs or HFMs and predicted their structures with AlphaFold2 (see Methods). Structures with SFMs show a similar structure to the wildtype with an *α*CTD conformation (Fig. 6C) while the ones containing the set of HFMs show a conformational change similar to *β*CTD (Fig. 6D). The latter suggests that destabilizing the interface between NTD and *α*CTD could induce the CTD metamorphic behavior of RfaH CTD.

**Fig. 6.**
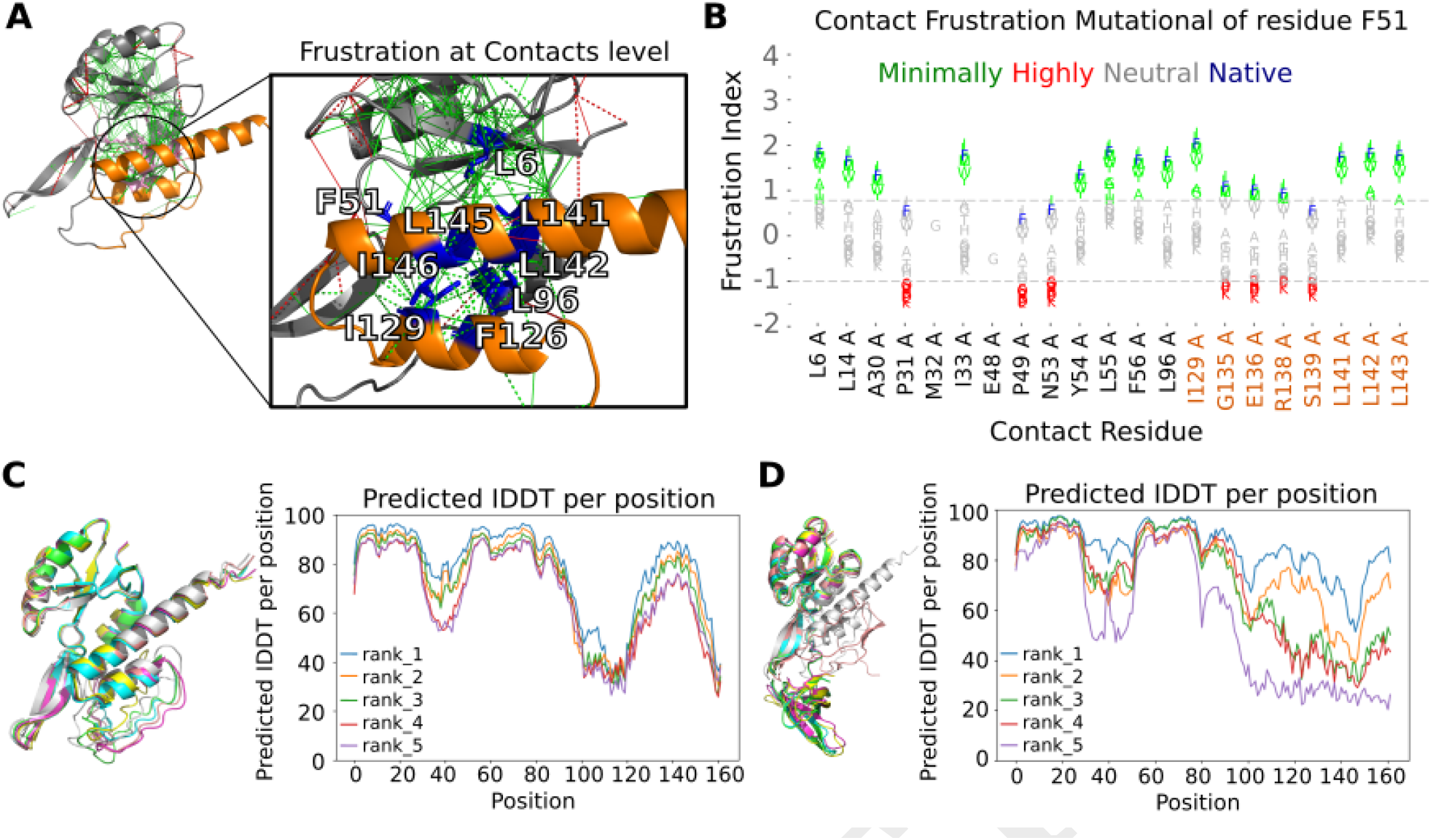
Frustration analysis of a metamorphic protein conformational change. **A)** FrustraEvo Mutational index results. Red lines correspond to highly frustrated interactions and green lines to minimally frustrated interactions (see Methods). Orange backbone corresponds to the interdomain region (CTD) and residues in blue and sticks are the 9 interface residues. **B)** Frustration changes upon mutation for F51 using FrustratometeR. The x-axis shows the residues with which the residue, either wild-type (Phe) or mutated, establishes contacts in the structure. In the y-axis we show the mutational frustration index for the contacts. The wild-type amino acid identity is shown in blue and the variants are coloured according to their frustration state. **C)** AlphaFold2 top 5 predicted models superimposed for RfaH containing different sets of mutations for SFMs and **D)** HFMs.

The previous analysis might not reflect realistic changes that could be introduced by nature. Therefore we investigated which sequence changes were introduced by evolution when comparing RfaH to NusG, its non-metamorphic homolog. Six of the 9 interdomain residues change their identity in the RfaH/NusG sequence alignment (i.e. NusG-like mutations: L6V, I129V, L141V, L145I, I146F and L142S, Fig. S13A). We introduced these mutations into the RfaH sequence and found that 4 out of 5 of the top AlphaFold2 structure predictions display a *β*CTD-like fold (Fig. S14A). The local frustration profiles of these mutations (Fig. S13B) shows that 4 of them are SFMs (L6V, I129V, L141V, L145I), one changes the frustration values from minimal to neutral (I146F) and only one (L142S) is an HFM. When L142S alone is introduced in RfaH, AlphaFold2 returns 1 model with an *α*CTD fold, 2 with loop-like CTD structures and 2 with a *β*CTD fold (Fig. S14A). From the 115 residues that change their identity in the RfaH/NusG sequence alignment, L142S is the only case where the *α*CTD changes to adopt a *β*CTD-like conformation (Fig. S14B) upon AlphaFold2 structure prediction. The CTD conformation adopted by the L142S mutant is less similar to the one of NusG when compared to that obtained when the 6 NusG-like mutations are introduced. Therefore, it seems that not only is the introduction of frustration necessary to trigger a conformational change in RfaH but also there is a need to fine tune the minimally frustrated contacts in the interdomain region.

This example illustrates how conserved frustration patterns can be used to generate hypothesis-driven experiments to modify specific biophysical properties in the context of protein engineering strategies.

## Discussion

We have introduced the analysis of energetic patterns across protein families based on the conservation of frustration levels. These patterns reveal physicochemical constraints related to stability and function, providing a biophysical interpretation of the impact of sequence divergence over evolutionary timescales.

We have shown that the analysis of conserved frustration patterns in individual protein families leads to the identification of functional constraints that correlate with changes either in stability (PDZ, SH3 and KRAS, Fig. 2C, 2E and 2G), function (PDZ, SH3 and KRAS, Fig. S2) or structural conformation (RfaH, Fig. 6). The comparison of frustration patterns across protein families points to regions of functional diversity (Hemoglobins, Fig. 3, RAS family Fig. 4 and SARS2 Fig. 5A and 5B) and specificity (SARS-CoV-2, Fig. 5C). In the case of the SH3 and PDZ domains, FrustIC values of minimally frustrated residues correlate with experimental ddPCA fitness values (Fig. 2C, and 2E). For the metamorphic RfaH protein, the analysis of conserved frustrated contacts led to the identification of key residues involved in conformational transitions. The best example is Leu142, proposed to play an important role in holding the metamorphic domain from transitioning from the all-*α* to the all-*β* conformation. The SH3 and RfaH examples highlight the importance of the conservation of frustration in an evolutionary context to identify meaningful functional constraints beyond what can be seen by sequence conservation alone.

In the case of the closely related *α* and *β* globin families that constitute the hemoglobin molecule, the differential roles in the transport of oxygen and contribution to the assembly of the quaternary complex is translated into a significant difference in the number and location of energetically conserved positions corresponding to protein-protein interaction sites and salt bridges. In contrast, for more diverse protein families, like the RAS superfamily, in which the sequence divergence signal might start to vanish, it is still possible to gather insights from frustration patterns that helps us go beyond sequence information for characterizing functional differences. We found out that the local contacts-based frustration patterns, but not the single residue frustration patterns, are conserved among the RAS, RHO, RAB and RAF subfamilies despite the large evolutionary distances between members. Some specific residues-residue interactions are systematically frustrated (mostly involving Asp57 and Lys117) in all families showing strong evolutionary pressure to maintain those conflicts despite sequence and functional divergence. This is expected since these residues are part of the active site and, therefore, associated to the common GTP-binding function of the RAS superfamily. Additionally, SDPs found in these protein families define differentially conserved specificity sites, whose interpretation in functional terms is neither easy nor direct. In these cases, the analysis of frustration conservation provides an additional layer of information. Finally, we have shown how a comparative analysis of frustration conservation can be performed in an unsupervised and automatic manner, leading to the identification of potential functional adaptations of protein families. As an example, we performed an analysis for 22 Coronavirus protein families. Despite an observable correlation between average FrustIC and SeqIC across protein families, the relationship between these two quantities is modulated by different aspects such as the phylogenetic diversity of the family or their propensity to protein disorder. By taking these factors into account and as shown with the PLPro example, frustration conservation analysis can be used for identifying positions that are likely relevant either for stability or function. Analysis of frustration changes across phylogenetic trees of emergent pathogens can identify novel adaptations in them as well as to mark residues to be taken into consideration when studying novel variants The evolution of protein families is constrained by both a narrow margin of stability and foldability energetics in the context of demanding functional requirements. In these margins, too many minimally frustrated regions might hinder functional evolution while the presence of too many highly frustrated regions will prevent folding from happening (39). Frustration conservation analysis within protein families can be used to define the theoretical limits to preserve function throughout evolution revealing the interplay between sequence, structure(s), dynamic and function(s) (40). Now that high quality protein structure models can be obtained for the members of any protein family, our frustration conservation analysis strategy stands as a valuable tool to increase the level of functional annotation in biological databases (41). Moreover, we envision that energetic local frustration, both at the level of single proteins or families, could be used as a novel measurement to complement the training of state-of-the-art machine learning methods such as protein language models as a way to improve their predictive/generative capabilities. Finally, the examples shown here illustrate how the approach we present can guide applications such as the construction of artificial proteins or the prediction of the risk of new natural pathogenic protein variants.

## Methods

### Local frustration

The frustration index, FI, allows one to localize and quantify local energetic frustration in protein structures (10, 42). Given a pair of contacting residues, their interaction energy is compared to the energies that would be found by placing different residues in the same native location (mutational frustration index) or by creating a different environment for the interacting pair (configurational frustration index). When comparing the native energy to the energy distribution resulting from these decoys, the native contacts are classified as highly, neutrally or minimally frustrated according to how distant the native energy is from the mean value of the energy of the decoys, taking into account the standard deviation of the distribution. An analogous approach can be used to calculate the FI for single residues (single residue frustration index, SRFI). In this case, the set of decoys is constructed by shuffling the identity of only one residue, keeping all other parameters and neighboring residues in the native location, and evaluating the total energy change upon mutation, i.e integrating the interactions that the residue establishes with all its neighbors. The configurational and mutational pairwise contacts FIs
have the following thresholds to define the different frustration states for the interactions, as proposed by Ferreiro et al. (10, 42): if FI*<*=-1 then the interaction is highly frustrated. If FI*>*=0.78 then the interaction is minimally frustrated. If -1*<*FI*<*0.78 then the interaction is neutral. In the case of the SRFI thresholds for single residues, if SRFI*<*=-1 then the interaction is highly frustrated. If SRFI*>*=0.55 then the interaction is minimally frustrated. If -1*<*SRFI*<*0.55 then the interaction is neutral.

### FrustraEvo’s pipeline

FrustraEvo calculates how conserved local frustration is in a set of protein structures that belong to the same protein family. It can be used in two different modes depending on which local frustration index is processed: 1) Single residue mode: using the SRFI, 2) Contacts mode: using either the configurational or the mutational pairwise contacts indices.

The input consists of 1) a MSA in FASTA format with sequences composed solely by the standard 20 amino acids code (other characters accepted by FASTA are replaced by a gap), and 2) A set of protein structures (experimental or models) in PDB format corresponding to the same set of sequences contained in the MSA. The ID of each protein sequence within the MSA should match its corresponding structure file name (without the .pdb extension). Sequences within the MSA should match exactly the ones contained in the PDB files.

Sequence and Frustration information content, SeqIC and FrustIC respectively, are calculated using information theory concepts. SeqIC is calculated from aligned residues in the MSA, based on the distribution of amino acid identities. It is calculated from the MSA using the ggseqlogo R package (43).

FrustIC is calculated for aligned residues in a MSA or equivalent contacts across proteins in the MSA, based on the frustration states mapped into the residues from the structures. A reference protein is selected to define over which residues or contacts the conservation calculations are calculated. The reference structure can be defined by the user or otherwise FrustraEvo selects the protein that maximizes the sequence coverage of the MSA.

In FrustraEvo’s single residue mode, the Frustration Information Content (FrustIC) for each column in the MSA is calculated based on the the Shannon entropy formula:

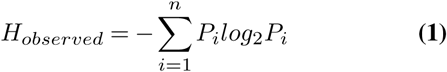

where *P*_*i*_ is the probability that the system is in frustration state i. The probabilities are normalized such as 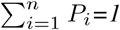, where M is the number of possible frustration states. For the frustration index, we consider minimally, neutral or highly frustrated states, therefore, M=3. To take into account background probabilities, the information content is calculated as *FrustIC =H*_*max*_ *- H*_*observed*_. Generally, it is considered that Hmax is reached for a uniform distribution of states: then 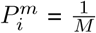 and *H*_*max*_ *= log*_2_*(M)*. Nevertheless, if states are not equally likely to occur a background probability distribution of states should be used to estimate Hmax. We used the distribution of states reported by Ferreiro et al (10) as background frequencies to calculate the Hmax for FrustIC calculations (minimally frustrated 0.4, highly frustrated 0.1 and neutral 0.5).

Similarly to the single residue mode, in FrustraEvo’s contacts mode, a reference structure is taken to define the contacts to be evaluated. Taking in consideration that the MSA was ungapped according to the reference structure, FrustraEvo calculates the frequency of having a contact between columns i, j in the MSA, on each structure in the dataset. Where i, j ∈ [1, N], with N being the number of columns in the ungapped MSA. As a result FrustraEvo will calculate, for each possible contact, according to pairs of columns within the ungapped MSA, the information content contributions from each frustration state. The FrustIC of a given contact will be calculated as the sum of the individual contributions from each frustration state. The background frequencies are the same as for the single residue mode. Plots for the frustration and sequence logos and contact maps are made using the ggplot2 R package.

### Data Visualization

We have developed the Multiple Sequence Frustration Alignment (MSFA) visualization to compare FrustratrometeR results across multiple protein sequences (e.g. Fig 1A). This type of plot consists of a heatmap for which each cell contains the residues in the MSA. Each cell is coloured according to their SRFI in the corresponding structures, i.e., minimally frustrated residues are colored in shades of green, neutral in gray and highly frustrated in red.

In addition to the previous, we designed the consensus MSFA to summarize and visually compare the FrustIC and SeqIC values across multiple protein families. In this case, each cell depicts the consensus sequence of the MSA of each protein family and the letter size is proportional to its SeqIC value. The cells’ background color corresponds, in shades from green through gray to red, to the median frustration value of that residue across all structures in the family. The background of the cell is white when FrustIC*<*0.5.

### AlphaFold2 Structure Predictions

Unless mentioned, for all analysis the 3D structure of each sequence contained in the final MSAs was determined using AlphaFold2 with default parameters. The best ranked model (rank1) was considered.

### PDZ, SH3 and KRAS workflow / data

To analyze GRB2-SH3 and PSD95-PDZ3 protein domains and KRAS with FrustraEvo, we generated MSA alignments for each family. To this extent, the sequences of reference, chain A from PDB 2VWF in the case of GRB2-SH3, chain A from PDB 1BE9 in the case of PSD95-PDZ3 and P01116-2 for KRAS, were blasted against the non-redundant NCBI protein database (from April 2021) using Blast v2.11.0 with default parameters. Hits with e-value ≥ 0.05, query coverage*<*70% and hits containing the words “artificial”, “fragment”, “low quality”, “partial”, “synthetic” were filtered out. The remaining hits were clustered using CD-Hit (44) with default parameters. Representative sequences were then aligned using MAFFT v7.453 (45) in the case of the SH3 and PDZ families and using HMMer’s hmmalign method in the case of KRAS, both with default parameters. The resulting MSAs contained 173, 679 and 2500 sequences for GRB2-SH3, PSD95-PDZ3 and KRAS respectively. For each of the sequences, AlphaFold2 models were produced. Their mean pLDDT values considering ronly residues that are contained in the alignment region of the reference structure were 89.3, 78.3 and 81.6 for SH3, PDZ3 and KRAS respectively. Finally, the MSAs and the AlphaFold2 models were analyzed with FrustraEvo.To calculate the correlation between experimentally determined stability and binding scores and FrustraEvo results, we used the ddPCA fitness scores from Faure et al. (15). From the supplementary files provided, we used: “JD_PDZ_NM2_bindingPCA_dimsum128_filtered_fitness _replicates.RData” and “JD_PDZ_NM_dimsum128_filtered _fitness_replicates.RData” to extract the results for PSD95-PDZ3 “JD_GRB2_epPCA_bindingPCA_dimsum128_fitness _replicates.RData” and “JD_GRB2_NM2_stabilityPCA _dimsum128_fitness_replicates.RData” to extract the results for GRB2-SH3.

In all cases, when loading the data files in R, only the dataframe “singles” was used in our analysis. We modified the numbering of PSD95-PDZ3 dataframes (column “Pos”) to start in 15 and end in 98 instead of 1 and 84 to match the real position of the domain in the protein sequence of reference (chain A from PDB 1BE9). For each position of both protein domains, the mean fitness value for each protein position is considered in the comparison to SeqIC and FrustIC. For the KRAS example, fitness values were obtained from the Supplementary table 4 (sheet 2) from the work of Weng et al. (16) and the mean fitness value was calculated for position and type of assay (AbundancePCA and all BindingPCA assays). Finally, we calculated the Pearson correlation between the mean fitness stability and binding data of GRB2-SH3, PSD95-PDZ3 and KRAS individually and their corresponding FrustIC or SeqIC values computed with FrustraEvo.

### Hemoglobins workflow / data

We have retrieved all non-redundant mammalian hemoglobins (n=21) present in PDB (by April 2022; Table S6),splitted them into two non-redundant structure sets of *α*- and *β*-globins and calculated their frustration patterns using FrustratometeR (14). Three MSAs were built, containing: all the *α*- and *β*-globins together, only *α*-globins, and only *β*-globins, the last two subsetted from the first one. Finally, for each MSA, we computed SeqIC and FrustIC values using FrustraEvo single residue mode.

### RAS superfamily workflow /data

A total of 160 human protein sequence IDs of the RAS superfamily were extracted based on Figure 3 from the work of Rojas et al. (17). Nine proteins were not included in the analysis as they were no longer present in Uniprot or their IDs did not match any entry. The sequences are grouped into ARF (n=26), RAB (n=64), RAN (n=1), RAS (n=38) and RHO (n=22). Since the RAN family only contains one human sequence, it was discarded from our analysis. Sequences were aligned with MAFTT v7.453 (45) with default parameters and their models were generated with AlphaFold2 (mean pLLDT scores are ARF=86.3, RAB=77.4. RAS=83.5, RHO=82.6).

### Unsupervised analysis of the SARS-CoV-2 and related coronaviruses proteins

We retrieved all homologues to proteins within the SARS-CoV-2 proteome with known sequences across coronaviruses (see following subsections) and generated models with AlphaFold2. After filtering out those proteins for which the structural models did not have enough quality, we processed a total of 22 protein families (Table S3). We applied the S3Det software (5) to subdivide the set of proteins into subfamilies and find their SDPs. Finally, we used FrustraEvo to obtain SeqIC and FrustIC values for each subfamily. The different steps of the pipeline are explained in more detail in the next sections.

### Coronaviruses homologs retrieval and MSAs building

Reference sequences for all SARS-CoV-2 proteins were retrieved from the reference genome MN985325 according to the annotations of the genbank file from the supplementary data provided in the work of Gordon et al. (46) (strain USA-WA1, file called “2020-03-04719B-Gordon et al 2020 - SARS-CoV-2 USA-WA1 Genome Annotation.gb”), with the following correction in line 408 “13442..16236” changed for “join(13442..13468, 13468..16236)”. Then each nucleotide reference sequence was translated and blasted against the non-redundant NCBI protein database (from March 2021) using Blast v2.11.047 with default parameters, a maximum of 100000 hits and e-value*<*0.05, excluding taxids 2697049, 2724902, 2724903, 2724904 which correspond to SARS-CoV-2 related sequences and taxid 32630 which corresponds to artificial sequences. Afterwards, the hits that were not from the Coronaviridae family were also filtered out. Hits with less than 70% query coverage and hits containing the words “artificial”, “fragment”, “low quality”, “partial”, “synthetic”, “Severe respiratory syndrome coronavirus 2” or “SARS-CoV-2” in the description were also excluded. The remaining hits were aligned using MAFFT v7.453 (45) with default parameters. In the case of the non-structural proteins (nsp), some of the hits retrieved corresponded to the whole orf1ab, so the alignment was trimmed to the region containing each specific nsp according to the SARS-CoV-2 reference. Afterwards, the sequences were clustered using CD-Hit (44) with parameters -c 0.98 -s 0.90 and the representative sequences were aligned with MAFFT.

### Classification into subfamilies and SDPs detection

The software S3Det (5) was used to classify the protein datasets into subfamilies and detect SDPs for each viral protein, following the same steps as in (47). Table S2 specifies the number of clusters determined per viral protein as well as the depth and length of each protein MSA.

### Structural Modeling and models quality filtering

Structure models for all proteins contained in the MSAs were generated using AlphaFold2 (8) with default parameters. To ensure high quality of the models, each MSA was trimmed to fit the length of a PDB of reference of SARS-CoV-2 proteins. When more than one PDB structure was available, the one that maximized the MSA coverage was selected. The reference PDBs for each MSA and the trimming positions are in Table S3. Transmembrane proteins M, nsp4 and nsp6 did not have a PDB available and were removed from the analysis. Finally, only high quality models, i.e.: mean pLDDT per SDP subfamily ≥ 80, were considered in the frustration analysis (Table S3 and Fig. S6).

### Phylogenetic balance

We calculated the phylogenetic diversity of each S3Det cluster of the 22 families of viral proteins (Fig. S7). The phylogenetic balance is represented by the standard deviation of the subgenus classes proportions across the cluster. When the standard deviation is low, it means that the cluster is balanced in terms of phylogenetic variability, i.e.: if there are 2 or more subgenus classes present, they are in similar proportions. On the contrary, when the standard deviation is high, the cluster is unbalanced, i.e.: the cluster is mostly represented by one of the subgenus classes.

### The PLpro example

A total of 122 non-redundant PLpro coronavirus homologous sequences were obtained and divided into 4 subfamilies which coincided with the following subgenera in Betacoronavirus: Sarbecovirus (n=31), Nobecovirus (n=11), Merbecovirus (n=35) and Embecovirus (n=45) (Table S3). In addition, and by making use of the extensive amount of experimental structures that are available for SARS-CoV-2, we retrieved all deposited PLpro structures in the PDB (n=29) and used them as an additional dataset that we also processed with FrustraEvo. The rationale behind this is that analyzing local frustration patterns across multiple structures from the same protein allows us to study the energetic determinants of the protein taking into consideration the conformational diversity of its native state. Flexible regions that exist in various frustration states across different structures will not show a high conservation signal while those that do not vary much will.

### Metamorphic protein workflow/data. Orthologs selection

While *Escherichia coli* RfaH (UniProtID: P0AFW0) is the only protein with available structures for both metamorphic folds, further proteins were selected as metamorphic RfaH orthologs based on the following criteria from bibliography. Four orthologs from *Salmonella typhimurium* (sequence identity: 88%), *Klebsiella pneumoniae* (80%), *Vibrio cholerae* (64%) and *Yersinia enterocolitica* (43%) have been demonstrated to be able to substitute *E. coli* RfaH function in vivo (48). Additionally, deleterious mutation of RfaH in *Y. pestis* and Y. *pseudotuberculosis* exhibits lipopolysaccharide defects similar to *E. coli δ*rfaH and therefore metamorphic behavior (48). Therefore, all protein sequences for these RfaH orthologs were retrieved for further analysis.

Selecting metamorphic orthologs for RfaH is not a straight-forward task as experimental confirmation is lacking for the majority of RfaH related sequences. We first retrieved all sequences from the IPR010215 entry in the InterPro database (49) and clustered them at 90% identity using CD-HIT (50) which gave us a total of 1004 sequences. As a strategy to determine which sequences are likely to be metamorphic homologs for RfaH in *E. coli* we followed these criteria: 1) for each RfaH sequence, there must be at least one reported NusG sequence in the same organism in the Uniprot database (51), 2) the full-length RfaH protein sequence must be predicted to fold into the autoinhibited, *α*-folded C-terminal domain (CTD) structure for RfaH (PDB 5OND); and 3) the isolated CTD of RfaH must be predicted to fold into the canonical *β*CTD fold (PDB 2LCL, 2JVV). The CTD was considered to start from the first residue forming secondary structure in the PDB 2LCL (residue 115, pattern KVII). We randomly selected sequences from the 1004 total set of entries in the redundancy reduced IPR010215 set until we completed 30 proteins that fulfilled the above mentioned criteria. In order to assure a high quality set of potentially metamorphic homologs, we manually confirmed that each instance to be added to the analysis fulfilled the mentioned criteria.

Finally, AlphaFold2 models (mean pLDDT=71.97 ± 3.8) were generated for each sequence in the ortholog set; min pLDDT=65.1, max pLDDT=79.4). It is worth to mention that metamorphic proteins use to contain regions with lower pLDDT (linkers and the metamorphic domain) scores due to their conformational diversity. For this reason we considered all structures without applying any further quality filter based on their mean pLDDT score.

### Interface residues identification

To detect those residues potentially involved in the fold-switch (the 9 interdomain residues, we selected those residues that: 1) establish interdomain residues contacts between the two domains according to the contact maps obtained with Frustratometer and further processed by FrustraEvo; 2) are located in the metamorphic region and their FrustIC*>*0.5 based on the contacts mode; 3) are present in more than 50% of the analyzed models; 4) have more than 50% of their interactions minimally frustrated; and 5) have at least 3 minimally frustrated interactions with other CTD residues. A total of 9 interdomain residues satisfied the previous criteria: L6, F51, L96, F126, I129, L141, L142, L145, I146.

### SFM ad HFM *E. coli* RfaH mutant sequences

For each interdomain residue that was detected as mentioned before we used the FrustratometeR module to predict the frustration change upon mutating the native identity by all the other possible non native amino acids. For each residue we selected an amino acid identity that would maintain a frustration value as similar as possible to the native identity (similar frustration mutation, SFM) and another one that would introduce as much frustration as possible (highly frustrated mutation, HFM). We generated RfaH mutants containing both the set of 9 SFMs and the 9 HFMs.

### NusG-like RafH

We aligned the RfaH and NusG sequences from *E. coli* (Fig. S13A) and observed that 6 out of 9 of the previously identified interdomain residues have different identities between the two proteins. We generated a “NusG-like” RfaH sequence by replacing the 6 residue identities from NusG into RfaH.

### Computational infrastructure and software requirements

MN4, Minotauro, Power9. Plots were produced with ggplot2 and ggpubr R packages. In this project we have used the frustratometeR package to calculate the local frustration patterns for all the presented analysis (14).

## Code and data availability

FrustraEvo is implemented in Python 3 and R 4.1.2 programming languages and encapsulated in a Docker container (https://hub.docker.com/r/proteinphysiologylab/frustraevo). FrustraEvo code as well as all necessary data to reproduce the examples (input MSAs and protein structures) shown in this manuscript are available in: https://github.com/proteinphysiologylab/FrustraEvo. All the output data from FrustraEvo, for all examples, will be available in ZENODO upon publication and can be provided under request. It is also possible to generate the output data by using the FrustraEvo docker container.

## ACKNOWLEDGEMENTS

This work was supported by the VEIS project (001-P-001647) from the European Fund for Regional Development of the EU (2014-2020 ERDF Operational Program of Catalonia). RGP thanks Prof. Christine Orengo and Nicola Bordin for early discussions and data provision. AV is an ICREA professor. CP is a Juan de la Cierva Formación postdoctoral fellow. PGW is supported by the Center for Theoretical Biological Physics sponsored by the NSF grant PHY-2019745 and the D.R. Bullard-Welch Chair at the Rice University Grant C-0016.

## Notes

### Competing Interest Statement

The authors have declared no competing interest.

