## Supplementary Figures for "The Evolution of Local Energetic Frustration in Protein Families"

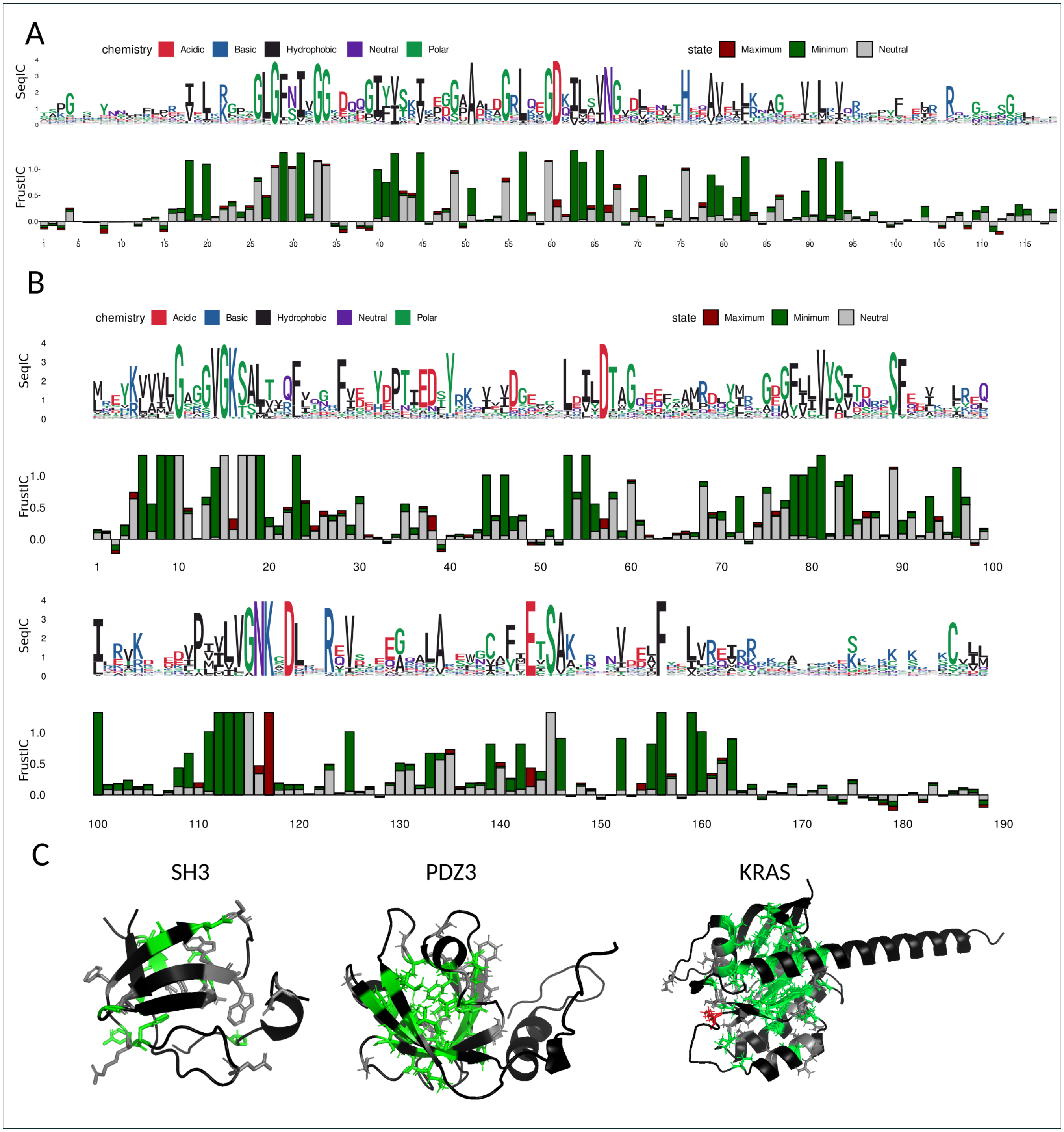


**Fig. S1.** **A)** Sequence and Frustration logo plots showing SeqIC and FrustIC values per MSA column respectively for PSD95-PDZ3. The numbering of the plot corresponds to the sequence of reference chain A from PDB 1BE9. Positions containing a gap in the sequences of reference are not considered in the plot. Protein PSD95-PDZ3 has been trimmed to match tested positions for mutation fitness in Rojas et. al. **B)** Sequence and Frustration logo plots for KRAS calculated with Rojas et al. data. Numbering corresponds to the protein of reference P01116-2. **C)** FrustIC results mapped to SH3, PDZ3 and KRAS proteins models. Residues with FrustIC$\leq$0.5 are shown in black. Residues with FrustIC$>$0.5 are coloured according to the frustration state that contributes more information to the overall FrustIC value.

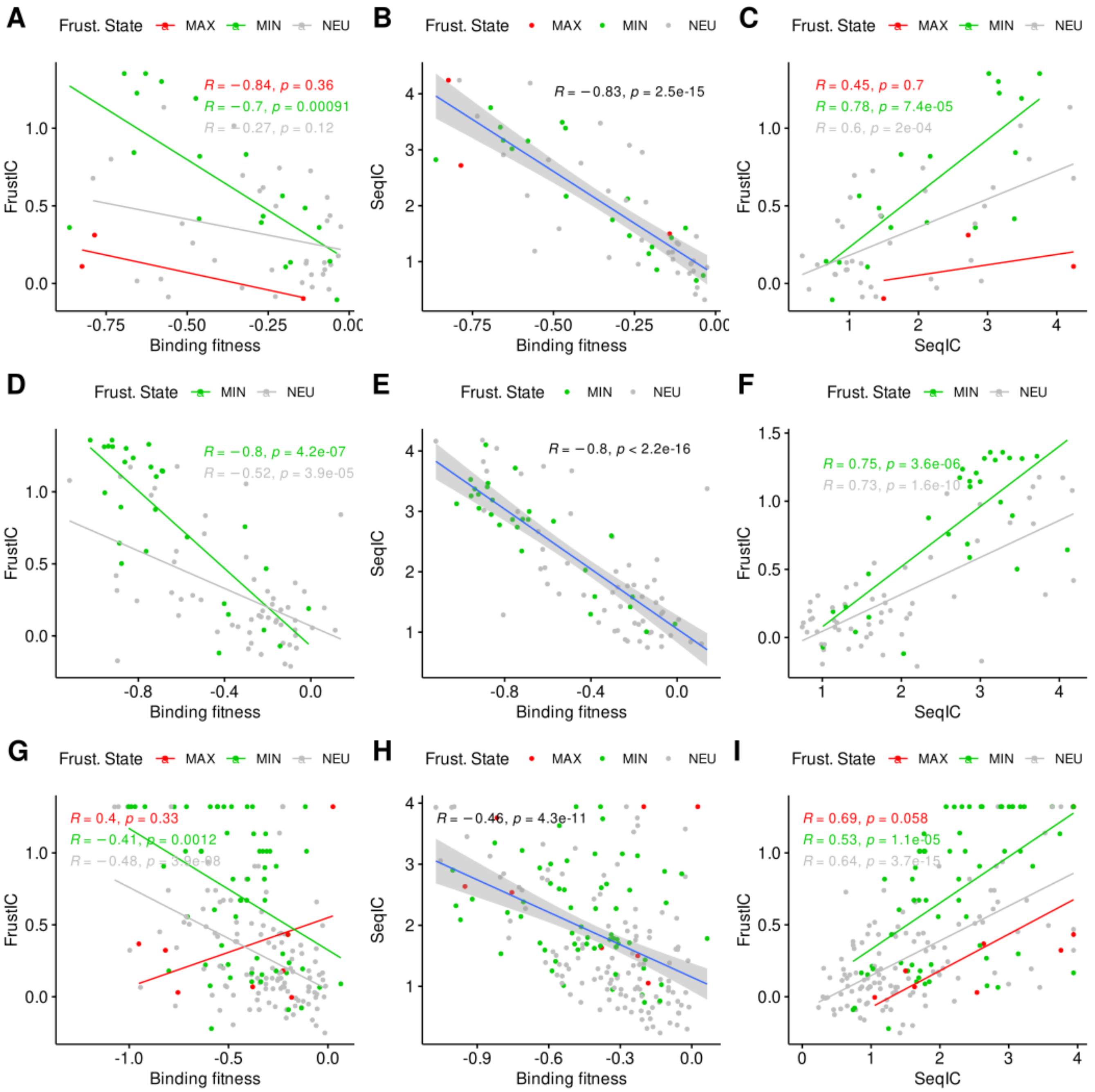


**Fig. S2. Correlation between FrustIC or SeqIC and binding fitness and FrustIC vs SeqIC in SH3 domain A, B, C)**, protein domain PDZ3 **D, E, F)** and KRAS protein **G, H, I)**.

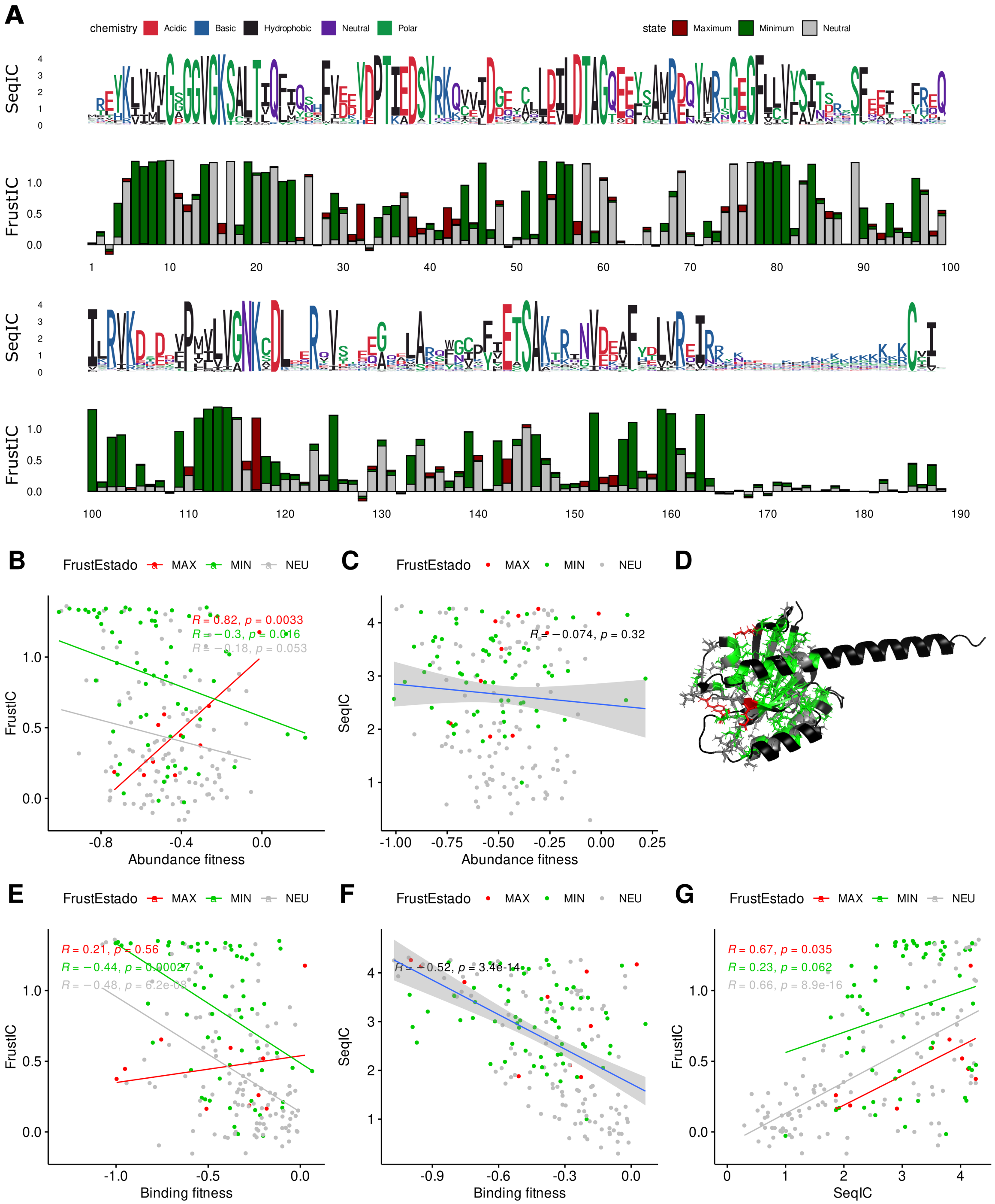


**Fig. S3. Results of KRAS with automatic retrieval of sequences of the protein family. A)** Sequence and Frustration logo plots for KRAS. Numbering corresponds to the protein of reference P01116-2. Correlation between FrustIC or SeqIC and abundance fitness **B and C)** or binding fitness **E and F**) and FrustIC vs SeqIC **G)**. **D)** FrustIC results mapped to KRAS proteins models. Residues with FrustIC≤0.5 are shown in black. Residues with FrustIC>0.5 are coloured according to the frustration state that contributes more information to the overall FrustIC value.

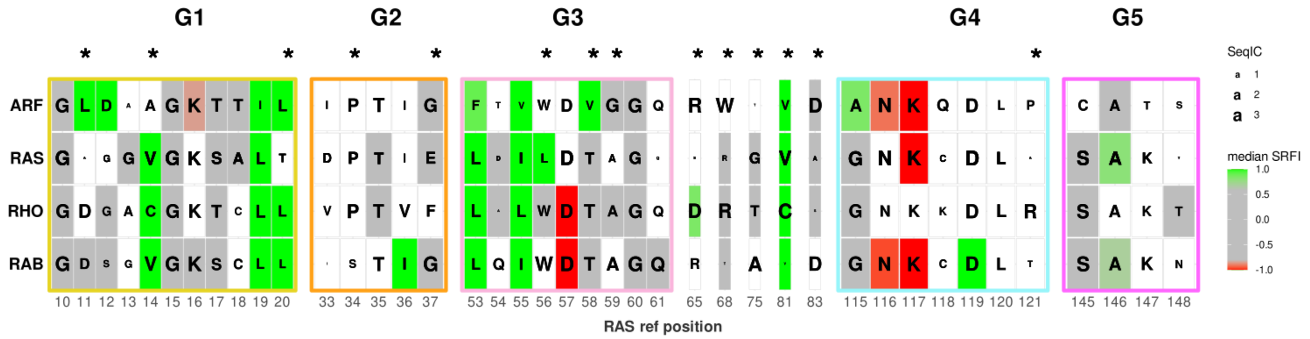


**Fig. S4.** Consensus multiple sequence frustration alignment (MSFA) comparing FrustIC and SeqIC results for the G1-G5 motifs and SDPs (marked with asterisks) in each of the subfamilies. Consensus amino acid identities are shown for each family. The size of the letter represents the SeqIC. The background color corresponds, in shades from green through gray to red, to the median single residue frustration index (SRFI) of that position across all structures in the family. White background means that FrustIC $\leq$ 0.5.

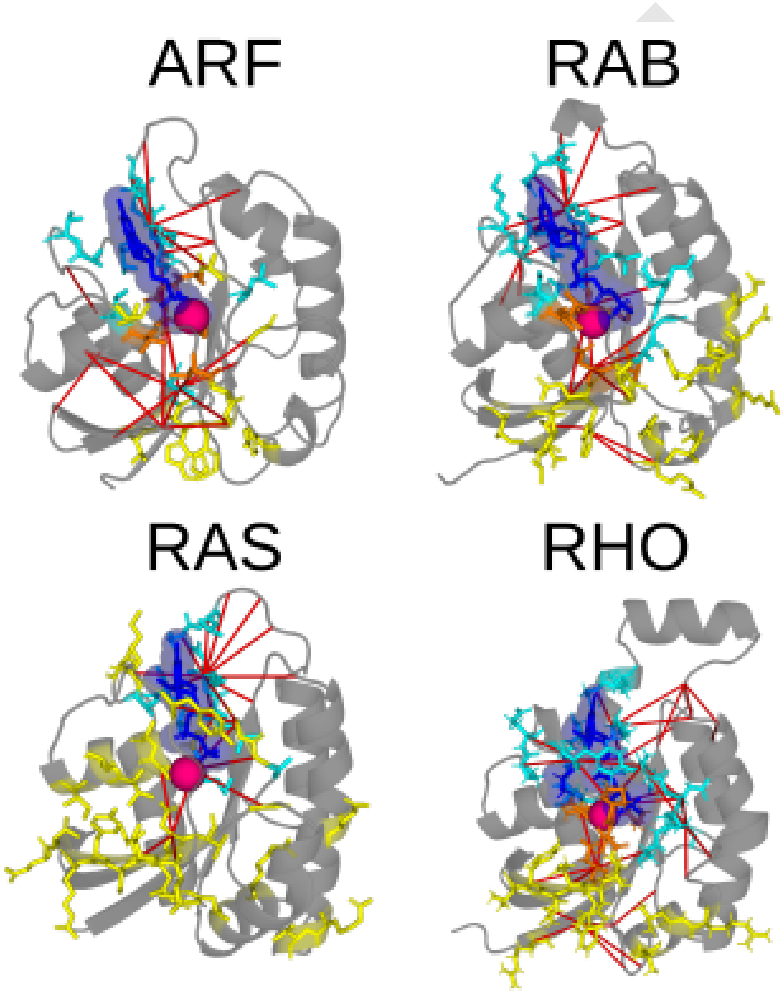


**Fig. S5.** Highly frustrated contacts between residues of the ARF, RAB, RAS or RHO proteins (PDB codes 7MGE, 3TKL, 1XD2 and 6BCB respectively) and the GTP nucleotide (in dark blue and sticks) or the Mg ion (in magenta and spherical shape). Additionally, residues colored in blue or yellow and represented in sticks are residues that interact with the GTP and Mg ligands or a protein partner included in the PDB file respectively (inter-residue distance $\leq 5$Å). When they are colored in orange means that they participate in both types of interfaces (protein and ligand).

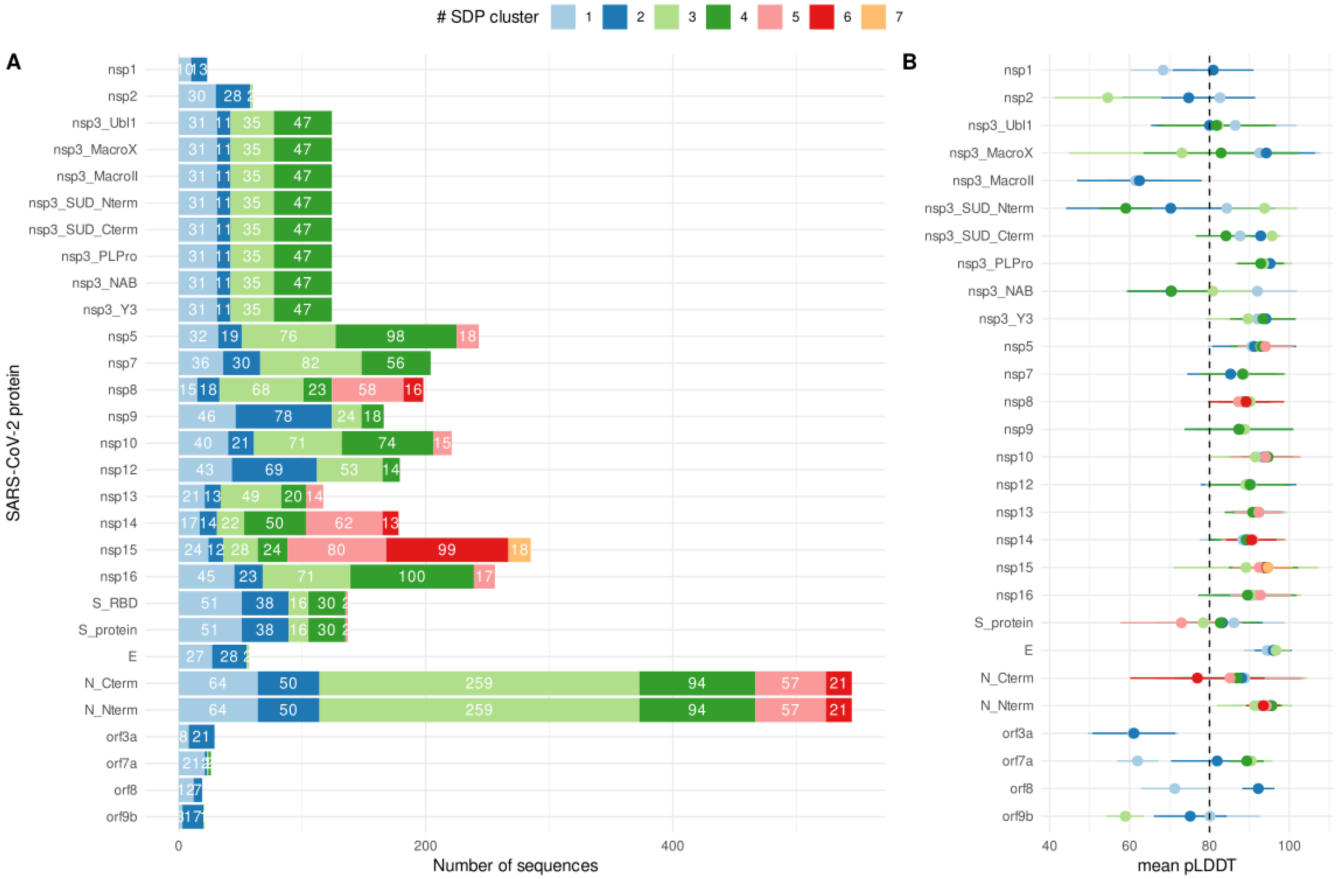


**Fig. S6. Coronavirus S3Det cluster and AlphaFold2 models quality. A)** Barplot depicting the distribution of sequences per S3Det cluster of all proteins or protein domains considered in the study (see Tables S2 and S3). Cluster with less than 10 sequences were not considered in our analyses. **B)** Mean pLDDT score per S3Det cluster of each protein or protein domain. The dashed line at pLDDT=80 represents the minimum quality threshold for a cluster to be considered. Below that we considered that the models are of low quality and therefore removed from the analysis.

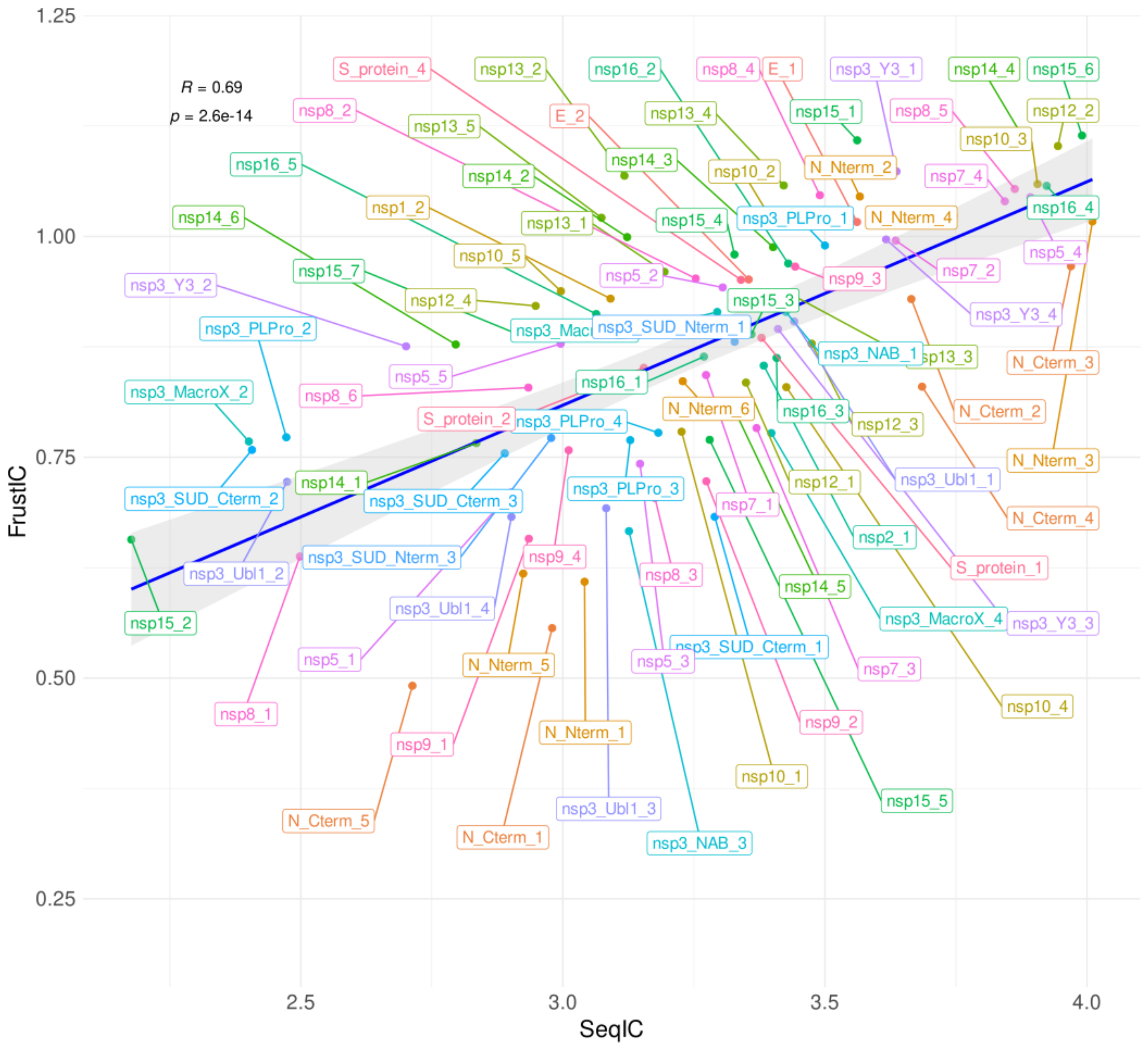


**Fig. S7.** Same correlation plot as in Fig. 5A showing mean FrustIC vs mean SeqIC per S3Det cluster computed for Coronavirus proteins (see Methods) but with all the data points labelled by protein and S3Det cluster.

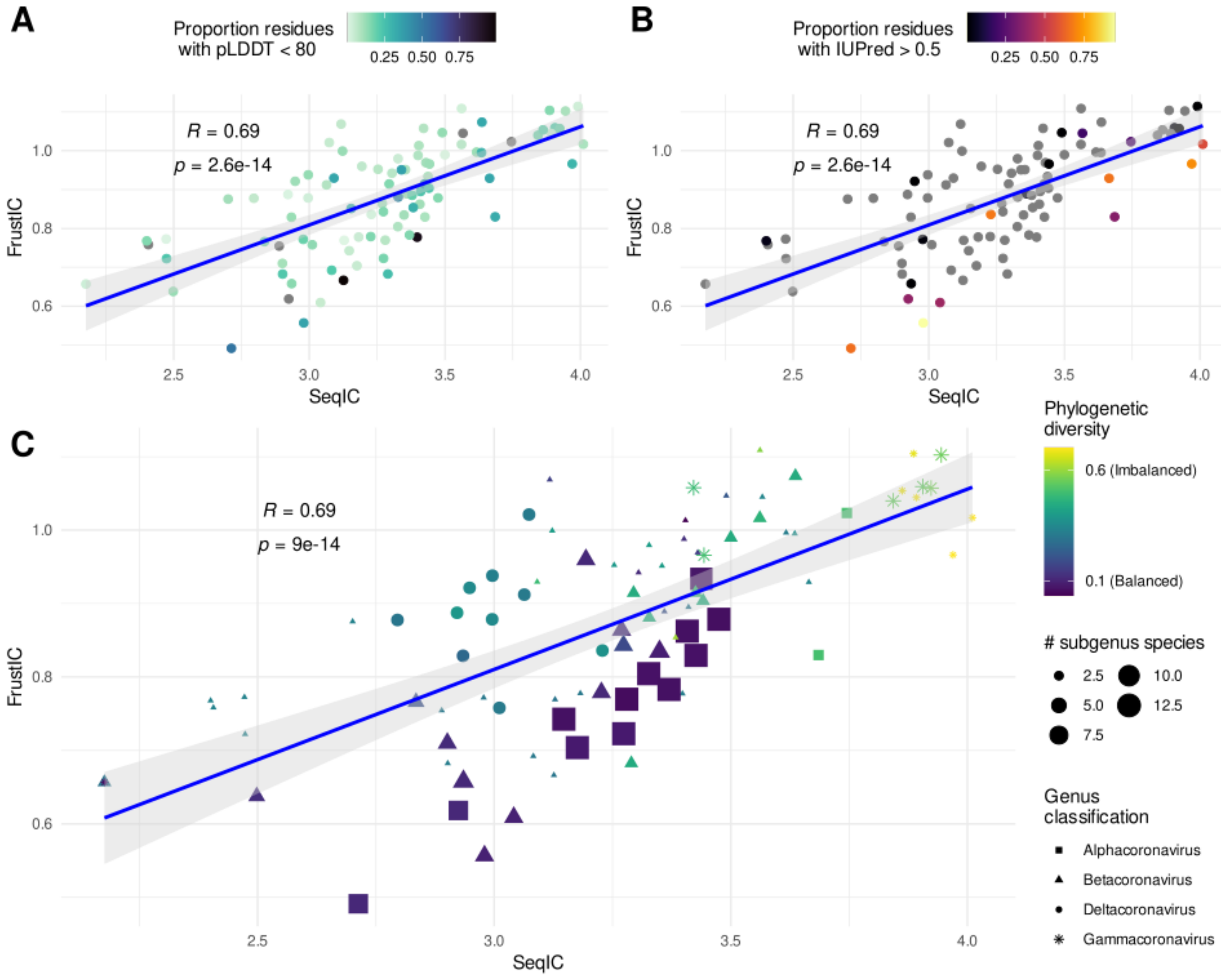


**Fig. S8. Factors affecting the expected correlation between SeqIC and FrustIC in Coronaviruses. A)** Quality of S3Det clusters models per protein represented by the mean proportion of residues with pLDDT $<$ 80 (low quality models). **B)** Disorder tendency of S3Det clusters models represented by the mean proportion of residues with IUPred $>$ 0.5 (disordered). Grey dots indicate that no residues were found with IUPred $>$ 0.5. **C)** Phylogenetic balance represented by the diversity of the subgenus classification within each protein and S3Det cluster (see Methods). The size of the point is represents the number of subgenus species represented by the considered sequences. The shape of the point refers to the corresponding genus classification (Alpha, Beta, Gamma or Deltacoronavirus) of all the sequences in each S3Det cluster.

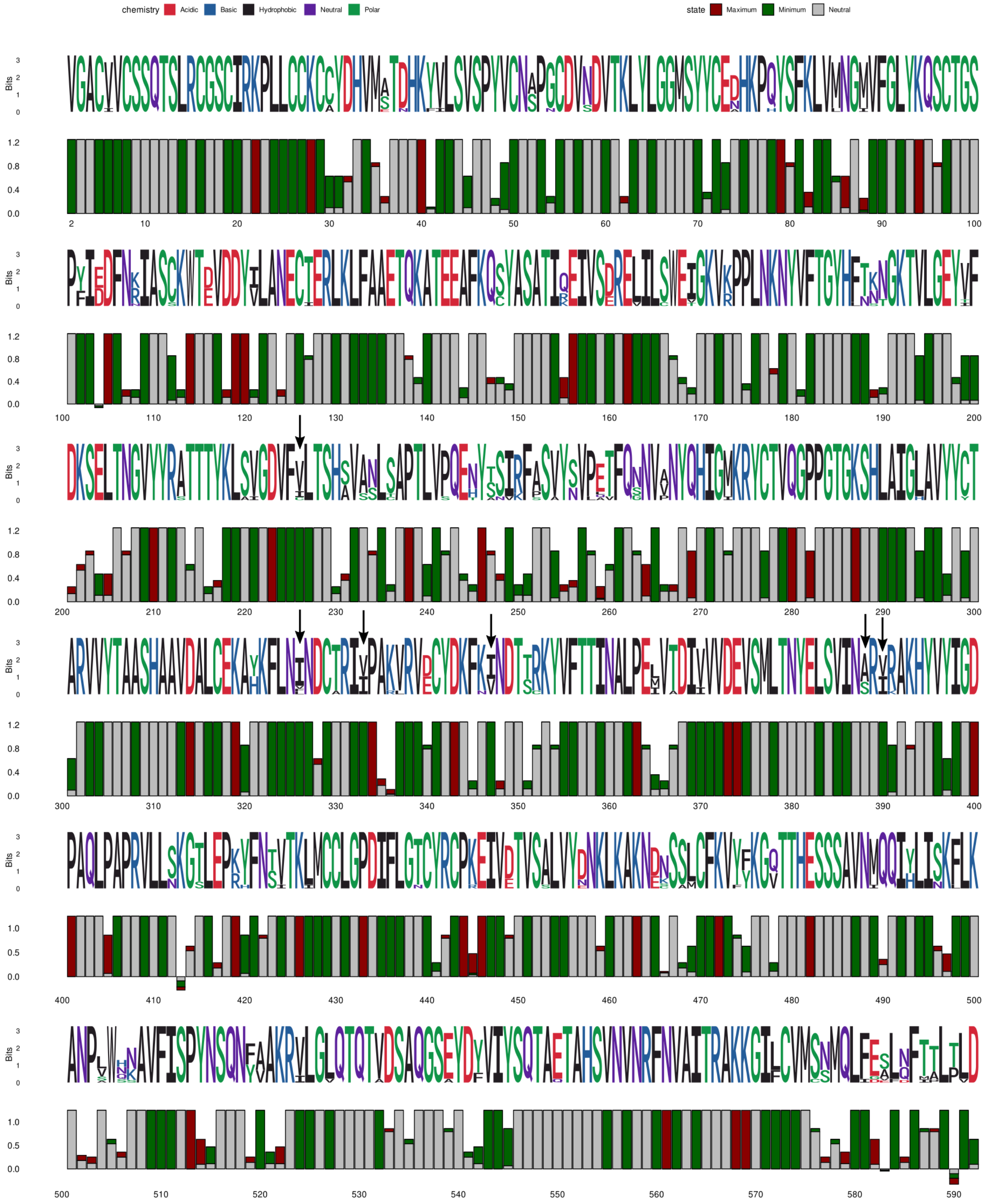

**Fig. S9.** Sequence and Frustration logo plots showing SeqIC and FrustIC values per MSA column respectively for the cluster 2 of Coronavirus protein nsp13. The numbering of the plot corresponds to the sequence of reference AYR18613.1. Positions containing a gap in the sequences of reference are not considered in the plot. Arrows point to examples of hydrophobic residues with lower conservation of SeqIC compared to FrustIC in a minimally frustration state.

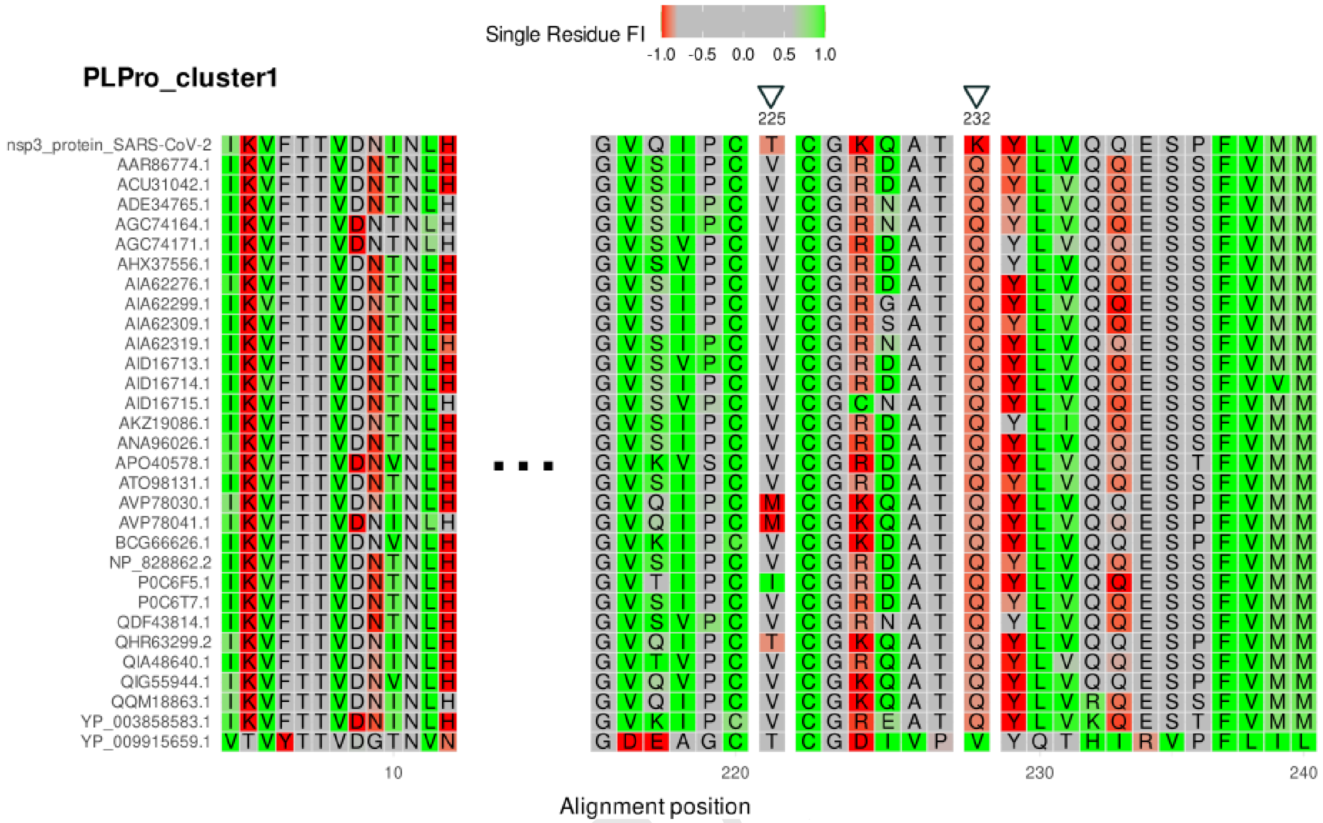


**Fig. S10.** Selected positions of the multiple sequence frustration alignment of PLPro S3Det cluster1 (Sarbecovirus). Highlighted positions 221/225 and 228/232 (Alignment position / SARS-CoV-2 numbering) representing interesting examples of change in frustration in the SARS-CoV-2 sequence (first row) compared to the rest of sequences.

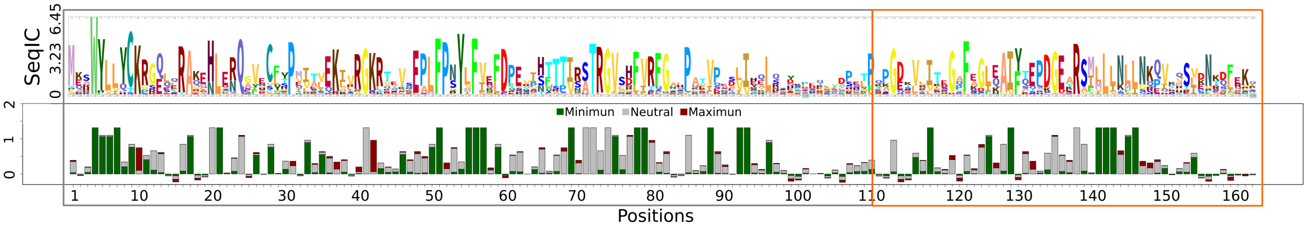


**Fig. S11.** Conservation of local frustration and sequence identity for RfaH family. FrustIC based on the single-residue level frustration index. In green are represented the minimally frustrated; in red highly frustrated contacts; and in gray neutral. In gray box is the non-metamorphic region and in orange box the metamorphic region.

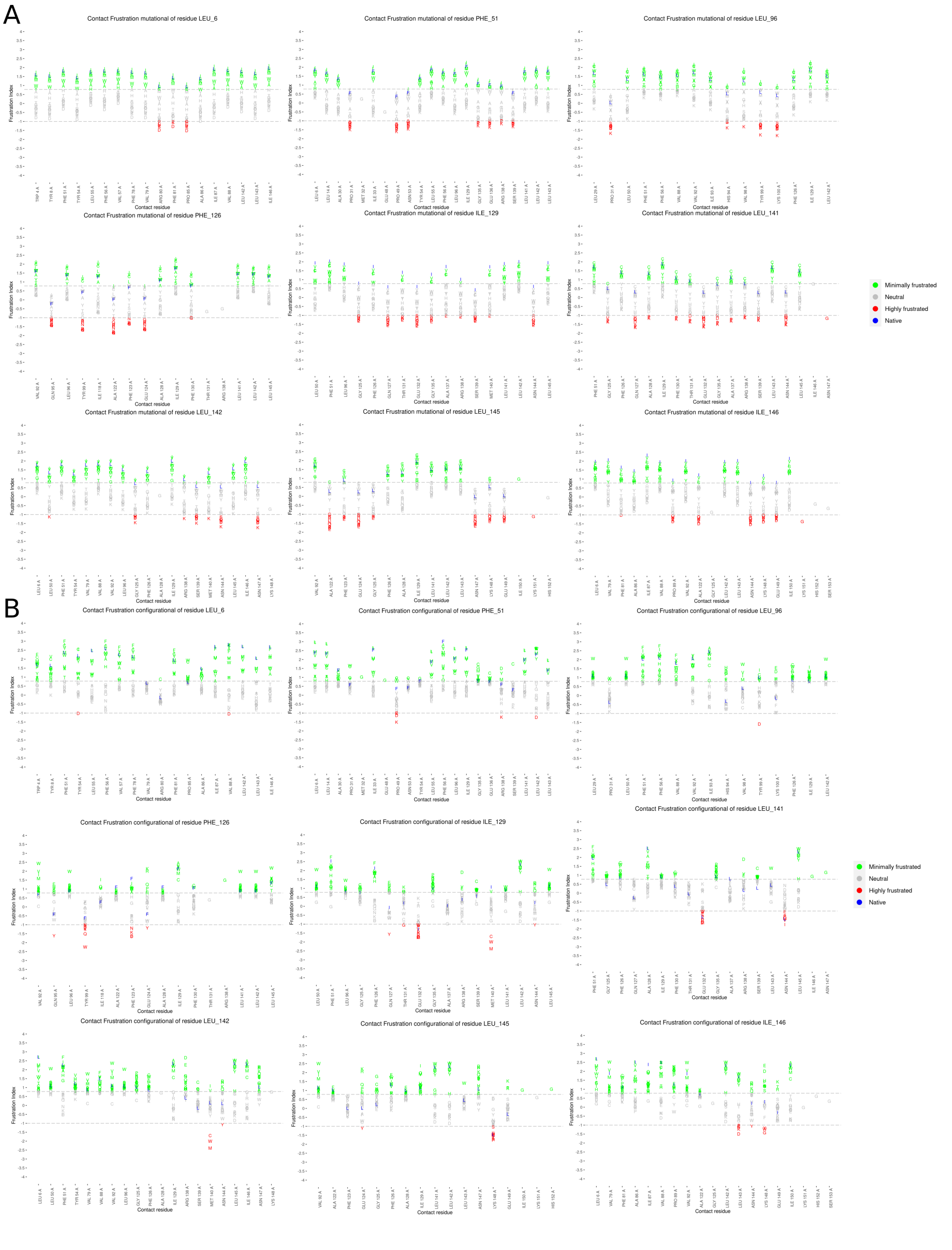


**Fig. S12.** Contact frustration changes, for A) mutational and B) configurational index, of a specific residue for all canonical amino acids alternatives. X axis: all possible contacts that form the native protein and the mutants are shown. Y axis: frustration values, the canonical amino acids alternatives are represented in letters and coloured based to their frustration value. Native variant appears in blue.

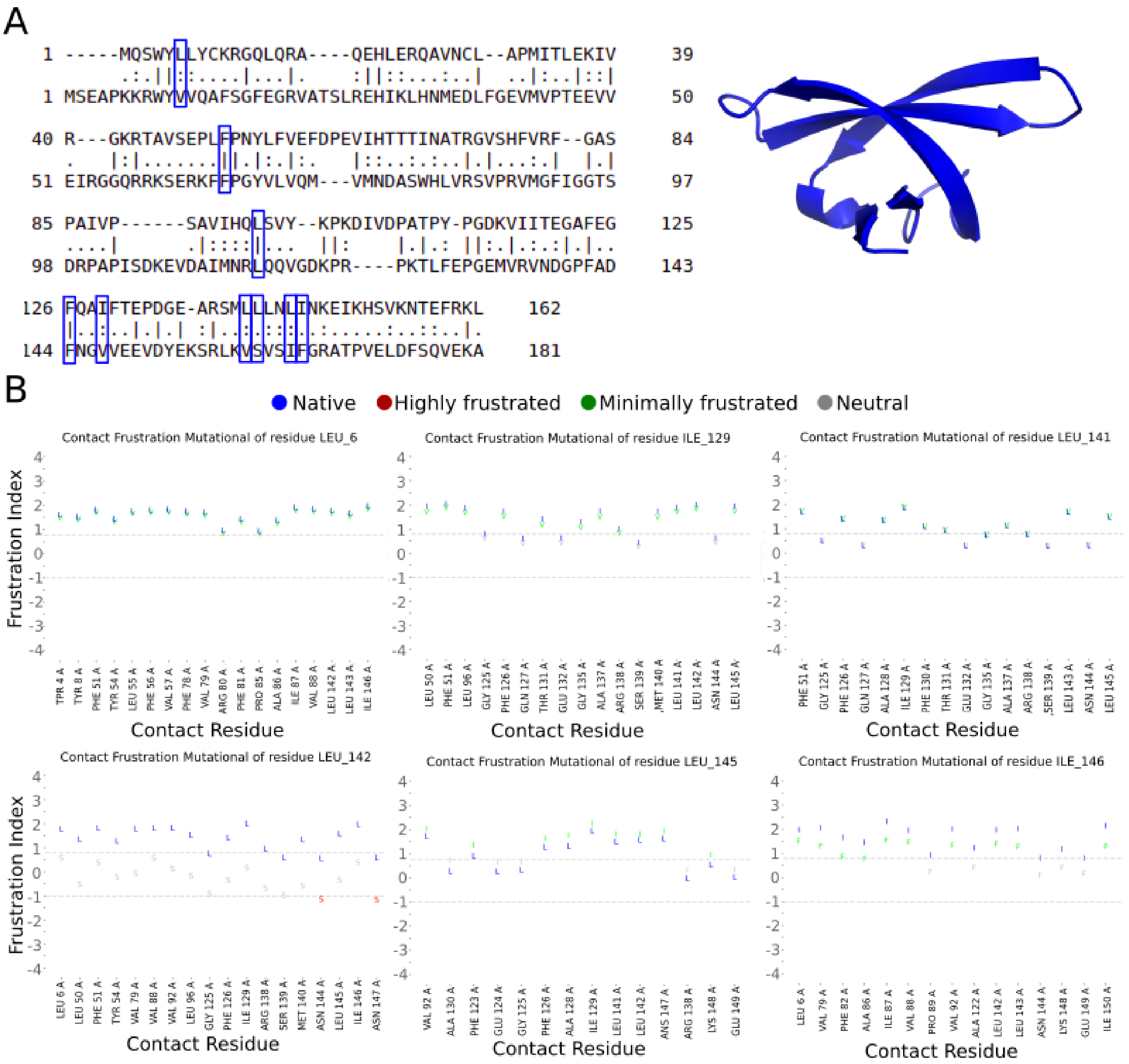


**Fig. S13.** A) Sequence alignment between RFAH and NUSG. B) Contact frustration changes for mutational index between native RFAH amino acids and NUSG equivalent positions. X axis: all possible contacts that form the native protein and the mutants are shown. Y axis: frustration values, the amino acids alternatives are represented in letters and coloured based to their frustration value. Native variant appears in blue.
